# Integration of distinct cortical inputs to primary and higher order inhibitory cells of the thalamus

**DOI:** 10.1101/2024.10.12.618039

**Authors:** Christian D. Puzzo, Rosa I. Martinez-Garcia, Hero Liu, Levi F. Dyson, William O. Gilbert, Scott J. Cruikshank

**Author notes:** These authors contributed equally to the work (Rosa I. Martinez-Garcia, Hero Liu).

## Abstract

The neocortex controls its own sensory input in part through top-down inhibitory mechanisms. Descending corticothalamic projections drive GABAergic neurons of the thalamic reticular nucleus (TRN), which govern thalamocortical cell activity via inhibition. Neurons in sensory TRN are organized into primary and higher order (HO) subpopulations, with separate intrathalamic connections and distinct genetic and functional properties. Here, we investigated top-down neocortical control over primary and HO neurons of somatosensory TRN. Projections from layer 6 of somatosensory cortex evoked stronger and more state-dependent activity in primary than in HO TRN, driven by more robust synaptic inputs and potent T-type calcium currents. However, HO TRN received additional, physiologically distinct, inputs from motor cortex and layer 5 of S1. Thus, in a departure from the canonical focused sensory layer 6 innervation characteristic of primary TRN, HO TRN integrates broadly from multiple corticothalamic systems, with unique state-dependence, extending the range of mechanisms for top-down control.

## Introduction

Top-down control of sensory and attentional processing by neocortical systems represents a landmark development in mammalian brain evolution^1–9^. The thalamic reticular nucleus (TRN), a GABAergic nucleus capable of gating the flow of thalamocortical information, is ideally positioned as a key mediator of top-down control. Axons descending from the neocortex to the thalamus (corticothalamic, “CT” axons) excite TRN cells via powerful collateral synapses. TRN cells, in turn, inhibit thalamocortical (TC) relay neurons, thus governing the strength and timing of signals that reach the cortex^10–16^. Consistent with its role as a cortical↔thalamic hub involved in sensation, perception, attentional processes, and arousal^16–33^, dysfunctions of the TRN have been linked to a range of neuropsychiatric conditions, including schizophrenia, Alzheimer’s disease, epilepsy, and sleep disorders^34–49^.

While the involvement of the TRN in top-down regulation has long been assumed, the precise mechanisms have been elusive. In part this is because many fundamental aspects of TRN circuitry, physiology, and principles of operation have been uncertain. However, a number of recent advances have been made, led by establishing the arrangement of TRN circuitry within the primary/higher order framework for thalamocortical organization^50–55^. This framework categorizes TC nuclei based on their major driving afferents. Primary TC cells are generally driven by ascending sensory inputs from the brainstem or periphery (“primary afferents”) whereas higher order (HO) TC cells tend to receive their strongest input from layer 5 of neocortex, after initial stages of processing (“secondary afferents”). It has become increasingly evident that the organization of TRN circuits is closely related to that of the TC nuclei, with each sensory sector of TRN divided into genetically, structurally, and physiologically distinct primary and higher order subdivisions^11,15,27,56–62^. Primary TRN neurons, situated in the central core of the nucleus, make synapses with primary sensory TC cells, and a surrounding shell of HO TRN neurons synapse with HO TC cells. The two TRN subtypes were found to have highly distinct physiological properties, both cell-intrinsic and synaptic. The former, which include robust differences in propensity for state-dependent bursting, suggest specialized integration and processing of synaptic inputs^23,27,58–60,62^

Despite the progress just described, significant gaps in knowledge remain. Among the most critical are uncertainties about cortical influences on the distinct TRN cell subtypes, with open questions ranging from basic anatomy of the cortical inputs to their functional consequences. For example, although it is generally agreed that the largest fraction of synapses in TRN arise from the neocortex^12,63,64^, the specific origins of those inputs are a matter of debate. Historically it has been assumed that cortical projections to the TRN originate almost exclusively from layer 6 (L6) CT cells of corresponding cortical areas, such as L6 of somatosensory cortex projecting to somatosensory TRN^10,11,16,50,65^. Important recent studies have challenged this assumption, reporting that L5 CT cells also project to TRN^66–68^ (and see ^32,69^). These cells can form morphologically large, “driver-like” synapses, contrasting with the small, “modulatory” synapses made by L6 CT cells^67–69^. If L5 projections to TRN are found to be prevalent, and are as functionally distinct from L6 projections as expected from CT studies involving thalamic relay neurons (reviewed in ^2,70,71^), it would significantly broaden the scope of possible top-down control mechanisms involving the TRN. However, the extent of L5→TRN projections remains controversial, with one research group indicating such projections are limited to frontal systems^68^, while another has reported that a wide range of cortical areas (both frontal and sensory) have L5 projections to the TRN^66,67^.

The L5 synapses from frontal areas to the TRN have been confirmed to be functional through physiological tests^68^, but those from sensory areas have not yet been functionally evaluated. In fact, little physiological information about CT influences on HO TRN exists at all, even for the more well-established L6 CT systems^21,28^. Given the dramatic differences in intrinsic neurophysiological properties of primary and HO TRN cells^23,27,59,60,62^, it seems likely that CT influences on their activities would be distinct, even if the properties of the CT synapses in the two systems were identical. At any rate, this hypothesis needs to be tested experimentally.

Considering that TRN is crucial to neocortical regulation of thalamocortical signaling, it becomes obvious that (1) identifying which CT cells and systems exert control over the primary and HO TRN, (2) clarifying their potentially unique synaptic properties, and (3) determining how those synaptic inputs are integrated to affect TRN cell activity (and resulting GABAergic output), will be essential to understanding the nature of neocortical top-down control. Here, we address these issues using anatomical, optogenetic and electrophysiological approaches.

We find that top-down synapses of the canonical L6 CT pathway from S1 cortex drive dramatically stronger spiking in primary than in HO cells of somatosensory TRN. This is due to both stronger synaptic input and more powerful T-type calcium currents in the primary TRN. The differences in T-type currents also cause distinct state-dependent response dynamics, with primary cell responses being phasic/depressing, and HO responses being relatively stable, when synaptic inputs arrive during hyperpolarized resting potentials. We also find that HO TRN cells receive additional, convergent, physiologically distinct inputs from layer 5 of S1 and motor cortex. Thus, in contrast to primary TRN, which receives input predominantly from the canonical layer 6 sensory CT pathway, HO TRN cells integrate a mixture of inputs from multiple CT pathways. This mixture mirrors the CT afferent patterns of the HO thalamocortical neurons to which they are linked. The stark differences in organizational and integrative properties among TRN cell subtypes are likely to have important implications for CT disynaptic inhibition and top-down regulation of primary and HO thalamic subcircuits.

## Results

### Part I. Distinct integration of sensory cortical L6 input by primary and higher order TRN neurons

Many thalamocortical nuclei receive convergent inputs from multiple CT systems with distinct properties (i.e., areal locations, laminar positions, synaptic morphologies and physiology)^2,5,71–81^. However, it has traditionally been thought that TRN neurons receive their CT inputs exclusively from layer 6 (L6), mainly from modality-aligned cortical areas (e.g., primary somatosensory cortex (S1) to somatosensory TRN)^10–12,16,50,65,82,83^. While that dogma is being challenged^66–69^, the importance of L6 CT projections to TRN is not in dispute. Therefore, we began our investigation of CT influences on somatosensory TRN subpopulations by examining the organization and impact of S1 L6 CT pathways.

### Anatomy of L6 CT projections

We first focused on the widely studied barrel cortex^84,85^, a subregion of S1 that processes vibrissae-related signals. Using Cre-dependent AAVs in NTSR1-Cre mice, we selectively expressed ChR2-eYFP in L6 CT cells^86–90^ of S1 barrel cortex (Figure 1A) and examined their projections to the somatosensory sector of TRN – the region of TRN that receives its thalamic input mainly from the VP and POm nuclei^11,15,19,27,56,60^.

**Figure 1.**
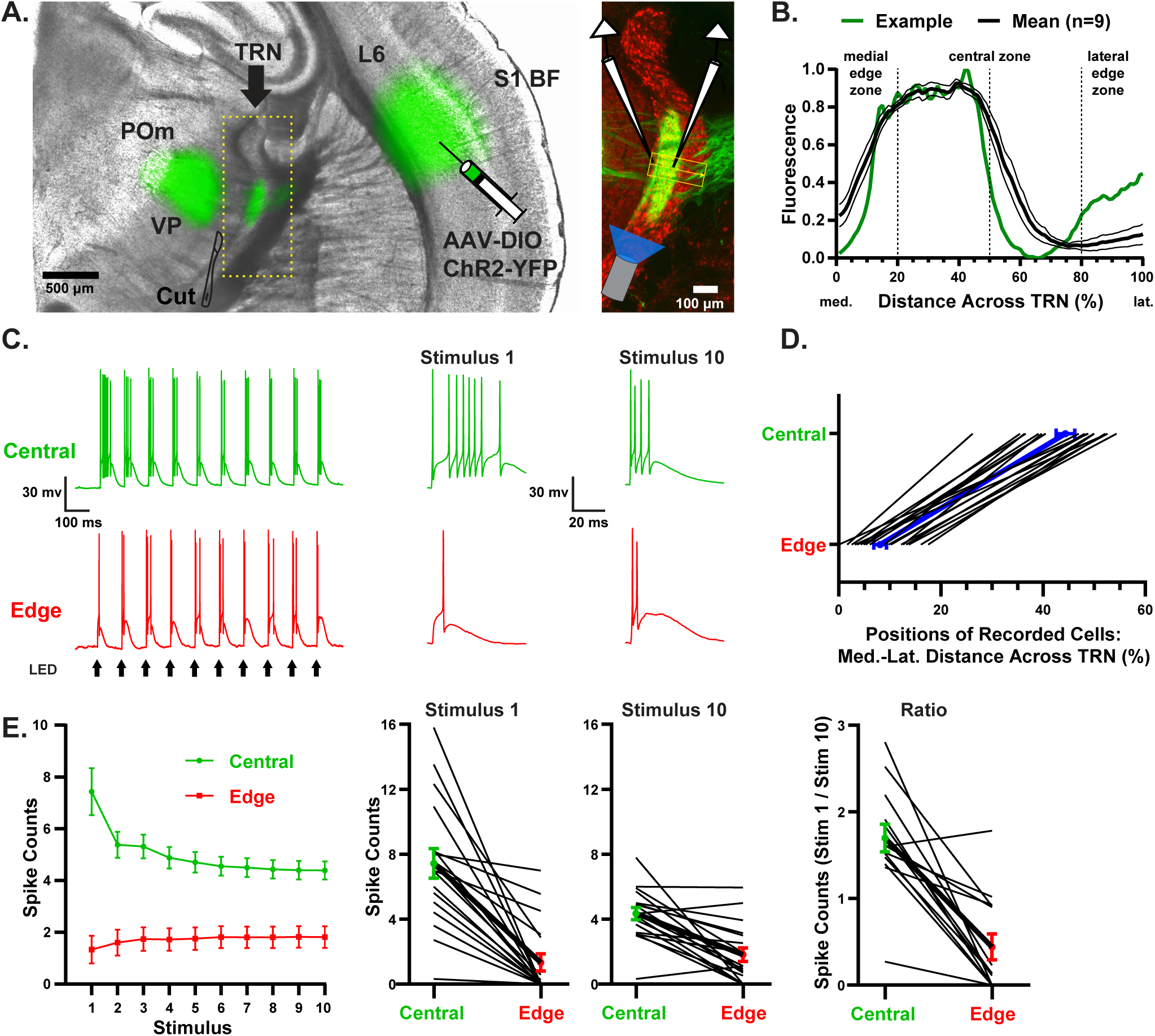
L6 CT input from S1 evokes distinct activity in primary and HO TRN neurons. (A) Experimental setup. **Left**, image of a live slice (300 μm) from an Ntsr1-Cre mouse injected in the barrel cortex with AAV1-DIO-ChR2-eYFP to induce ChR2-eYFP expression selectively in L6 CT cells. Image is an overlay of brightfield (gray) and eYFP (green). Before recording, the VP and POm thalamocortical nuclei were removed with a scalpel. ChR2-eYFP expressing CT axons were stimulated optically within the TRN and resulting postsynaptic responses were recorded in whole-cell mode from nearby TRN neurons in the central and medial edge zones of TRN (below). **Right**, magnified image of a 50 μm thick section from the same slice following immunohistochemistry for parvalbumin (red) and YFP (green) to identify TRN cells and L6 axonal projections, respectively. Schematic overlay: Paired whole-cell recordings from primary central and HO medial edge cells, with optogenetic stimulation of CT axons. TRN, thalamic reticular nucleus; POm, posterior medial nucleus; VP, ventral posterior nucleus; S1 BF, primary somatosensory cortex barrel field. (B) Fluorescent densities of L6 CT projections across the TRN. For each preparation, an ROI was drawn from the medial to lateral edge of TRN, intersecting the center of the projection (as illustrated in the right panel of A). Green line, the projection profile of the example shown in A. Black line, average profile ± SEM (n=9 preparations). The L6 CT projection densities in TRN are highest in the medial half of the central zone, and fall off sharply both medially and laterally. (C) **Left**, action potential activity evoked by optical stimulation of L6 CT axons (10Hz train of 1 ms pulses, 11.8 mW) in a pair of TRN neurons, one from the central zone (green) and the other from the edge zone (red). **Right**, responses to 1^st^ and 10^th^ stimuli in the trains (expanded time bases), illustrating burst-like responses selectively in the central cell. Spike counts from stimulus 1 to stimulus 10 decreased for the central cell and increased for the edge cell. Throughout the study, paired comparisons of central and edge TRN neurons were made from cell pairs recorded serially within the same slices, and baseline membrane potentials and holding potentials were kept at ∼-84 mV (= their approximate resting potentials)^60^ unless indicated otherwise. (D) Positions of recorded TRN cells (0 = medial edge of TRN). Thin black lines connect within-slice cell pairs (n=18 pairs). Thick blue line and symbols show means ± SEM. Edge cells were located within 20% of the medial→lateral distance across TRN (mean = 8.1 %). Central cells were between 25% and 55% of the medial→lateral distance across TRN (mean= 44.39 %). (E) **Far left**, group plot showing number of action potentials discharged by TRN cells for each stimulus in the 10 Hz trains (mean ± SEM; n= 18 cell pairs from 16 mice, 11.8 mW stimulus intensity). The spiking evoked by S1 L6 input was stronger in primary central cells than HO edge cells (mean spike counts across stim 1-10: Central = 50.0 ± 4.3, Edge = 17.2 ± 4.4; P<0.0001, n = 18 cell pairs, paired two-tailed *t*-test). **Middle**, numbers of action potentials evoked for stimulus 1 and stimulus 10 for all recorded cells (again lines connect within-slice cell pairs). More spikes were evoked in central cells than their paired edge cells for both stimuli (*p*<0.0003, paired two-tailed *t*-tests, Bonferroni corrections for multiple comparisons). **Far right**, Stimulus 1/Stimulus 10 spike count ratio for central/edge pairs. In primary TRN, the spiking evoked by S1 L6 input depressed during repetitive stimulation, while HO TRN spiking facilitated (Spike count ratio, Stim 1/Stim 10: Central = 1.7 ± 0.16, Edge = 0.44 ± 0.15; n=14 cell pairs from 13 mice, *p*<0.0001, paired two-tailed *t*-test).

We observed systematic non-uniform distributions of these projections to the somatosensory TRN. When AAV injections were made across the full thickness of L6, the peak of the TRN projection was focused ∼40% of the medial→lateral distance across the nucleus, corresponding to the primary central TRN zone defined previously by thalamic projections and molecular identity (see Introduction and Methods)^27,60^. There was also a lesser, but clear, projection to the higher order edge zone comprising the medial 20% of the somatosensory TRN. Labeling dropped off sharply in the lateral half of the primary central zone, and only increased slightly in the HO edge zone near the lateral boundary of TRN (Figure 1A-B)^80^.

Results from previous studies suggest that CT projections to the central and medial edge zones of TRN originate from distinct L6 sublaminae^80,83^. To test this here, we performed focal dual AAV injections in upper and lower L6 of NTSR1-Cre mice (using AAV-FLEX-ChR2-eYFP and AAV-FLEX-ChrimsonR-tdT), then made within-preparation comparisons of the corresponding thalamic targets (Figure S1A). In 17/18 mice the projections from lower L6 terminated more medially in the TRN than those from upper layer 6 (Figure S1B-C). This confirms previous conclusions^80,83^ and indicates that primary central and HO medial edge neurons of TRN receive input from largely separate groups of L6 CT cells – from upper and lower L6, respectively.

We next asked if the patterns of L6 CT projections observed for barrel cortex were repeated for non-trigeminal fields of S1 (e.g., the forelimb/hindlimb/trunk regions: “FL/HL/TR”), perhaps giving rise to convergent projections to common TRN targets. Thus, we expressed ChR2-EYFP in FL/HL/TR S1 L6 CT cells and compared their TRN projections to those from the barrel field. Within the central primary zone of TRN, projections from the distinct S1 fields to were clearly offset, with the FL/HL/TR CT axons terminating more laterally than those from the barrel field (Figure S1D-E)^83,91^. However, we also observed an additional moderate projection from FL/HL/TR to the HO medial edge zone of TRN that appeared to overlap with the HO TRN projection from the barrel field (Figure S1D-E). Taken together, these findings suggest that primary TRN neurons located in separate central subzones receive most of their L6 CT inputs from distinct S1 fields, but that HO TRN neurons located near the medial edge of TRN may receive overlapping/convergent CT inputs from multiple S1 fields.

### S1 L6 CT inputs drive dramatically distinct spiking activity in primary and higher order TRN

To examine the functional influences of the L6 CT projections, we next recorded activity evoked in primary and HO TRN cells by optogenetic stimulation of S1 L6 CT axons (10 Hz LED trains, 1 ms/pulse). For these experiments, ChR2 was induced in CT cells with Cre-dependent AAV injections across the full depth of barrel cortical L6 in NTSR1-Cre mice (Figure 1A).

Consistent with the anatomical results (Figure 1B), the spiking evoked by CT inputs in primary central TRN cells was far stronger than in HO edge cells (Figure 1C-E). Furthermore, there were striking differences in the dynamic patterns of the activity. Primary TRN cells responded to L6 inputs with robust, high frequency, spike bursts that progressively depressed during repetitive stimulation. In contrast, HO TRN cells rarely burst, and their spike rates generally underwent modest facilitation (rather than depression) to repeated input (Figure 1C-E).

### Differences in EPSC amplitudes evoked by L6 inputs contribute to the differences in spike rates among TRN subtypes, but dynamic patterns of the EPSCs are roughly similar across the TRN

We next sought to determine the underlying mechanisms of these cell type differences in CT-evoked spiking. We first tested possible roles of synaptic inputs carried by the L6 CT afferents (Figure S1) to the two classes of TRN cells, measuring the excitatory postsynaptic currents (EPSCs) evoked by L6 axon stimulation (recorded from the same TRN cells tested in Figure 1). If synaptic inputs were entirely responsible for the differences in spiking, we would expect to observe strong and depressing EPSCs in primary TRN, and weaker but facilitating EPSCs in HO TRN.

Indeed, EPSCs evoked in the primary central TRN were more than 2-fold stronger than those in HO edge neurons (Figure 2A-C; S2A), clearly contributing to their stronger CT-evoked spike rates. Note that these differences in EPSCs (as well as in evoked spike rates) were also observed when ChR2 expression in L6 CT cells was achieved by crossing Ntsr1-Cre with Ai32 mice, rather than through AAV FLEX methods (Figures 3A-F; S3). This indicates that the stronger responses in primary TRN cells are likely a function of the pathways per se, and not idiosyncrasies relating to the method of opsin expression – such as viral tropisms.

**Figure 2.**
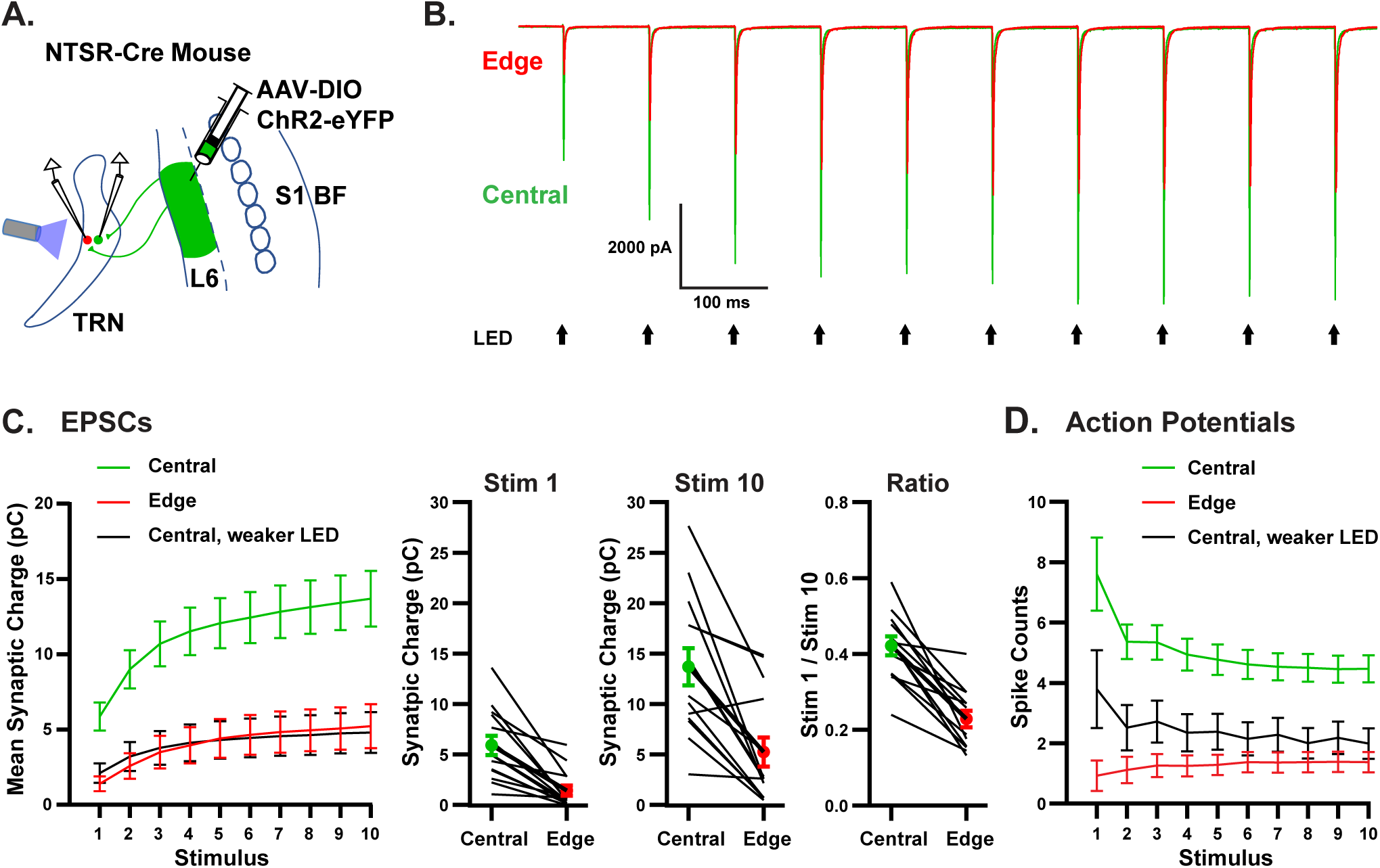
Differences in EPSCs evoked by S1 L6 input in primary and HO TRN: Influences on spiking. (A) Schematic of experimental setup. As for Figure 1, AAV1-DIO-ChR2-eYFP was injected into S1 barrel cortex of Ntsr1-Cre mice to induce ChR2-eYFP expression in L6 CT cells, then synaptic responses evoked by optical activation of the L6 CT projections were compared in pairs of central and edge TRN neurons. (B) Representative EPSCs from a cell pair evoked by matching optical stimuli (11.8 mW). Amplitudes of the EPSCs from the central cell (green) were nearly twice those from the edge cell (red). (C) **Far Left**, group plot of postsynaptic charge evoked by S1 L6 CT input for each stimulus of the 10 Hz trains in the central and edge neurons. The EPSCs were stronger in primary central cells (green) than HO edge cells (red) when stimulus intensities were matched for the two cell subtypes (11.8 mW CT stimuli; mean synaptically-evoked charge across stim 1-10: Central = 114.7± 15.8 pC, Edge = 40.6 ± 11.9 pC; *p*<0.0002, n = 14 cell pairs, paired two-tailed *t*-test). The black line shows the mean postsynaptic charge for the central cells when the intensity of the CT LED stimulus was lowered to evoke EPSCs of the same overall magnitude as those evoked in the edge cells when using 11.8 mW stimuli (mean reduced LED intensity = 3.79 ± 1.38 mW). **Middle,** both the 1^st^ and 10^th^ stimulus of the train evoked greater EPSCs in central cells than their paired edge cells (*p*<0.001, paired two-tailed Bonferroni *t*-test). **Right**, the Stimulus 1/Stimulus 10 EPSC charge ratios were below 0.6 for all cells, consistent with strong short-term facilitation across subtypes. However, the ratios were higher for central than edge cells, indicating statistically greater facilitation in HO edge cells despite the qualitative similarities (Stim1/Stim 10 EPSC charge ratios: Central = 0.421 ± 0.025, Edge = 0.228 ± 0.022; n=13 cell pairs from 11 mice, *p*<0.0001, paired two-tailed *t*-test). (D) To determine the contribution of the stronger CT-evoked EPSCs of the central cells to their greater spiking responses, the CT-evoked spiking in the central cells was tested at lower LED intensities – intensities at which central EPSC sizes matched those of their paired edge cells when evoked with 11.8 mW stimuli (mean EPSC plots shown in panel C). Lowering CT input strength this way reduced the evoked spiking of central cells (compare black vs. green plots), but not to the levels of edge cells (black vs. red), and spiking continued to depress slightly during repeated stimulation (Central weaker LED vs Edge 11.8 mW LED, *p*<0.03, n = 15 pairs, paired two-tailed *t*-test; Central weaker LED vs Central 11.8 mW LED, *p*<0.0005, n = 15 pairs, paired two-tailed *t*-test; Bonferroni corrections for multiple comparisons).

**Figure 3.**
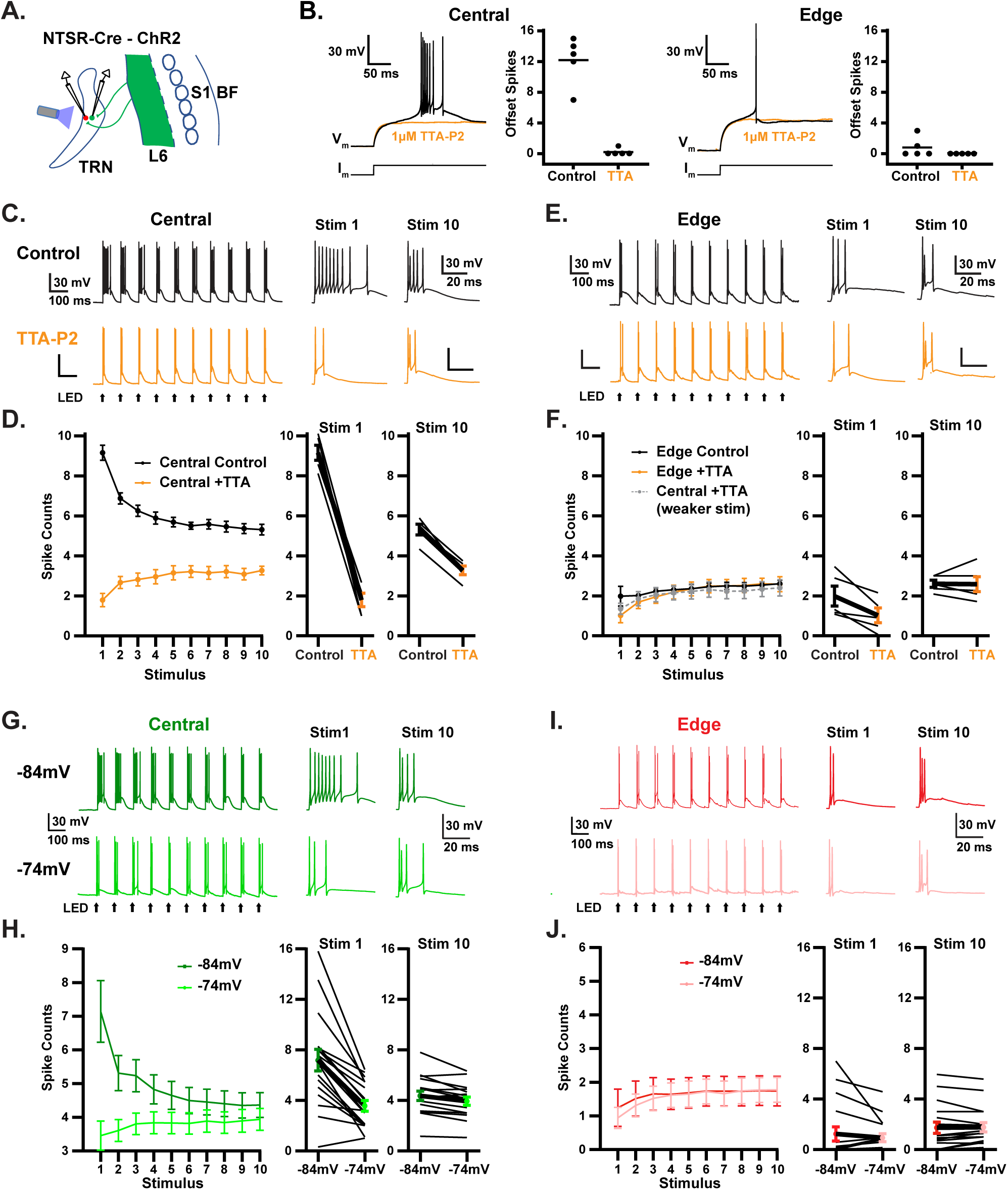
Voltage-dependent bursting mediated by T-type calcium channels enhance and shape responses to CT input predominantly in primary TRN. (A) Schematic of experimental setup. ChR2-eYFP was expressed in L6 CT neurons by crossing Ntsr1-Cre with Cre-dependent ChR2-eYFP reporter mice (Ai32; panels B-F), or by S1 injections of AAV1-DIO-ChR2-eYFP in Ntsr1-Cre or NTSR1-Cre x PV-Flp x Ai65F mice (panels G-J). Synaptic responses evoked by optical activation of the L6 CT projections were compared in pairs of central and edge TRN neurons before and during application of TTA-P2 (1 uM; a selective T-type calcium channel blocker; panels C-F), and during induced changes of the steady-state Vm (panels G-J). (B) **Left**, intrinsic bursting in a central TRN cell, and blockade of bursting during TTA-P2. “Offset bursts” were triggered by injecting negative current to reach ∼ –94 mV, followed by abrupt current removal. Group data shows number of spikes discharged within 150 ms after termination of the negative current. The 1 uM TTA-P2 eliminated bursting in all central cells. **Right**, same as left panels but for edge TRN cells. Edge cells had minimal offset firing in control conditions, and TTA-P2 blocked those responses. (C) L6 CT-evoked spiking in a central TRN cell before and during TTA-P2. **Top**, characteristic bursting responses to initial CT stimuli, with gradual reduction in spiking across the train. **Bottom**, application of TTA-P2 eliminated initial bursting and unmasked a facilitating spiking pattern. (D) **Left**, group plot showing the mean numbers of action potentials (± SEM) evoked by each stimulus before and during TTA-P2. **Right**, within-cell comparisons of the numbers of action potentials evoked by the 1^st^ and 10^th^ CT stimuli before vs. during TTA-P2 (n=5 pairs from 5 mice). (E-F) As in C-D except focused on edge cells (n=5 pairs from 5 mice, same slices as in C-D). Application of TTA-P2 had only minimal effects on edge cell responses. The gray plot (in F) shows central cell spiking when CT LED intensities were lowered to produce EPSC sizes matching those of their paired edge cells (edge cell EPSCs evoked by 23.5mW). When synaptic inputs were matched this way (and T-Type Ca++ channels blocked) central and edge spike responses were nearly identical (3F, gray vs. orange). (G) Effect of depolarization on L6 CT-evoked spiking in a central TRN cell. **Top**, spiking evoked from –84 mV showing initial bursting and decreased spiking by the end of the CT train. **Bottom**, depolarization of the steady-state Vm (to –74 mV) weakened responses, strongly reduced bursting, and led to facilitation. (H) **Left**, group plot showing the mean numbers of action potentials evoked by each stimulus in primary central cells. **Right**, within-cell comparisons of L6 CT evoked spiking for the 1st and 10th stimuli from – 84mV vs. –74mV steady-state potentials. Depolarization caused responses of central cells to weaken and develop facilitation, resembling the effects of TTA-P2 application in (D) (Mean spike counts: –84mV = 49.2 ± 4.5, –74mV = 37.9 ± 3.3; *p*<0.0001, n=17 cells from 15 mice, two-tailed paired t-test). Responses decreased for both Stim 1 and for Stim 10 (Stim 1: P< 0.0002; Stim 10: P< 0.006; paired two-tailed t-tests comparing –84 to –74 mV, n = 17 cells from 15 mice; corrected for multiple comparisons). However the suppression was stronger for Stim 1 than Stim 10, consistent with the switch from short-term depression to facilitation (P<0.0001; paired two-tailed t-tests comparing depolarization-induced decrease in evoked spike counts for Stim 1 vs. Stim 10, n = 17 cells from 15 mice). (I-J) Same as G-H but for HO edge cells. Depolarization had little effect on evoked spiking in edge cells, similar to the application of TTA-P2. Mean counts did not change significantly (–84mV = 16.4 ± 4.6, – 74mV = 15.7 ± 3.6; P=0.5980, n=17 cells from 15 mice, two-tailed paired t-test), nor did responses to Stim 1 or Stim 10 individually (Stim 1: P=0.7552; Stim 10: P=1; paired two-tailed t-tests comparing –84 to –74 mV, n = 17 cells from 15 mice; corrected for multiple comparisons).

To assess the degree to which the stronger EPSCs in the primary central cells contributed to their greater CT-evoked spiking, LED stimulus strengths were adjusted to evoke matching synaptic charge in the two TRN subtypes (within-preparation central/edge cell pairs; see Methods), and spike responses were tested using those stimuli. Under these conditions, the differences in CT-evoked spiking among TRN subtypes were clearly reduced, although they were not completely eliminated (Figure 2D). Thus, factors beyond the characteristics of the CT synaptic inputs alone must have contributed to the differences in spiking magnitude between the two TRN subtypes (below).

L6 CT EPSC in the two TRN cell subtypes had similar dynamic properties, with both undergoing strong short-term facilitation (Figures 2B-C, S2, S3), consistent with corticoreticular synapses described previously in unclassified TRN neurons^8,89,92–94^. This appears to require the calcium sensor, synaptotagmin 7, as deletion of the Syt7 gene virtually eliminated facilitation (Figure S4)^95–97^. Importantly, the facilitating EPSC patterns matched the dynamic CT-evoked spiking patterns of the HO TRN cells, but were opposite to the depressing spiking patterns of the primary TRN cells. Based on these mismatches in dynamics, and the previous analysis of magnitudes, it becomes clear that the differences in CT-evoked spiking among the two TRN subtypes must, at least in part, result from mechanisms downstream of the corticoreticular synapses. Thus, we next focused on possible roles of intrinsic physiological properties of the TRN cells.

### The TRN cell subtypes have different propensities for T-type calcium channel-dependent bursting, which differentially shape their spiking responses to CT synaptic input

We and others previously observed that primary and HO TRN neurons have distinct intrinsic physiological properties that could differentially influence their synaptic integration^23,27,59,60,62^; also see ^58,98–101^. One potentially powerful influence is the far greater propensity for bursting in the primary central cells. Unlike HO TRN cells, which exhibit minimal bursting, nearly all primary cells fire high frequency bursts of action potentials upon release from hyperpolarizing stimuli (Figure 3B)^27,59,60^. This firing behavior is consistent with stronger T-type (low-threshold) calcium currents^102–106^ in primary cells. In fact, here we found that blockade of the T-type channels by the selective blocker TTA-P2 completely eliminates such bursting (Figure 3B, S5)^107,108^.

We therefore asked if the greater intrinsic propensity for bursting in primary TRN cells contributes to their unique CT-evoked spiking. From resting potentials (∼ –84 mV), primary central cells responded to layer 6 CT input with high frequency barrages of action potentials that resemble intrinsic bursting; this synaptically-evoked bursting declined with repetitive activation, leading to the depressing phenotype (Figures 1C-E, 3A-D). We hypothesized that this pattern is largely driven by characteristics of T-type calcium currents, such that the progressive declines in spiking during repetitive synaptically-evoked activity result from the known voltage-dependent inactivation of those currents^102–104,106^. To test this, we applied TTA-P2 to block T-type channels, then measured responses to trains of L6 CT synaptic input (Figure 3C-F). Under such blockade, the differences in responses between subtypes were greatly reduced. The CT-evoked spiking in primary cells shifted from depressing to facilitating patterns, matching the patterns of HO cells (Figure 3C-F). Furthermore, the blockade of T-type channels reduced synaptically-evoked spike rates more in primary than in HO cells (Figure 3D-F; p < 0.015, paired t-test comparing % decrease in spike rates in central – edge cell pairs, 5 pairs from 5 mice), bringing the rates closer for the two subtypes.

Nevertheless, even in the presence of TTA-P2, a modest rate difference remained (Figure 3D vs. 3F), presumably due to the stronger synaptic inputs in the primary cells (Figure 2 above). Consistent with this interpretation, when CT stimulus intensities were adjusted so that the EPSCs of the primary and HO cells had matching magnitudes (still under T-Type Ca++ channel blockade), spike responses of the two cell subtypes became virtually identical (Figure 3F; p = 0.8701, paired t-test comparing central + TTA-P2 (weaker stim) vs. edge + TTA-P2, n = 5 cell pairs). Together these results indicate that the cell type differences in TRN spiking evoked by S1 L6 CT inputs are produced by a combination of greater synaptic input and stronger T-type calcium currents in the primary cells.

The cell type differences in T-type bursting may have important implications for state-dependence of CT modulation. Shifts in brain state during behavioral transitions, such as from quiescence to aroused wakefulness, can have profound effects on thalamic neurons^14,64,109–119^. For example, it often produces changes in firing modes between bursting and tonic activity, which is thought to depend on the voltage-dependence of T-type calcium channels. Consistent with this, depolarization of the steady-state V_m_ in primary TRN cells from their normal resting potentials (∼-84 mV) to –74 mV, shifts their evoked firing patterns from bursting to “tonic” modes^60^, due to T-type calcium channel inactivation.

Here we observed that such depolarization likewise transforms CT-evoked spiking of primary TRN cells to a less bursty, facilitating pattern (Figure 3G, H). In contrast, HO TRN cells have far weaker bursting overall (Figure 3B)^60^, and we observed that voltage shifts induced only modest effects on CT-evoked spiking in HO TRN, with relatively low spike rates and moderate facilitation occurring no matter the steady-state V_m_ (Figure 3I-J). These data suggest that changes in membrane potentials that accompany shifting brain states would have greater impact on activity and output patterns of primary than of HO TRN cells. Such differences could have important consequences for state-dependent inhibitory effects of the two TRN subcircuits.

### Part II. HO TRN cells integrate convergent top-down inputs from multiple CT pathways

An essential result from the preceding section is that the canonical CT pathway from L6 of S1 exerts far weaker effects on HO than on primary cells of the somatosensory TRN. Does this mean that top-down control over HO TRN is generally weak? An alternative possibility is that cortical influences on HO TRN are robust, but rather than originating predominantly from L6 of aligned cortical areas, they may emerge from other (perhaps multiple) CT pathways. To test this possibility, we examined the projections and synaptic effects of two additional CT pathways that have been associated with processing in somatosensory thalamus, the pathways from layer 5 of S1 and from M1^5,66,71,75,79,81^.

### L5 CT cells of S1 monosynaptically excite HO neurons of the somatosensory TRN

CT cells of layer 5 (L5) are known to make morphologically large, physiologically powerful, synapses with HO thalamocortical neurons (such as those in the POm) on their way to motor-related targets in the brainstem and/or spinal cord^65,71,73,74,82,120–126^. Classically, these L5 CT projections have been thought to bypass the TRN entirely^2,11,19,82^. While recent studies have challenged this assumption^66–68^ (also see ^32,69^), to our knowledge there has yet to be confirmation of functional synapses from L5 of sensory cortices to TRN.

Here we addressed these issues, applying optogenetic methods that allow selective excitation of L5 CT projections from S1, to examine possible synaptic effects in TRN. First, we expressed ChR2-eYFP in L5 CT neurons of S1 using stereotaxic injections of AAVs carrying Cre-dependent opsin genes into RBP4-Cre mice. In the neocortices of these mice, ChR2-eYFP expression was largely restricted to cells in layer 5 of S1 (Figure 4A), consistent with laminar patterns in prior reports^66,68,127,128^. Projections to thalamocortical nuclei were dense in POm, and significantly weaker in VPm, as expected for S1 L5 CT pathways (Figure 4B; reviewed above and in ^8^). In contrast with long-standing consensus, but in agreement with a recent anatomical study by Sherman and colleagues^67^, we observed projections to the TRN, with the highest densities along the outer edges of the nucleus – corresponding to HO TRN zones (Figure 4C).

**Figure 4.**
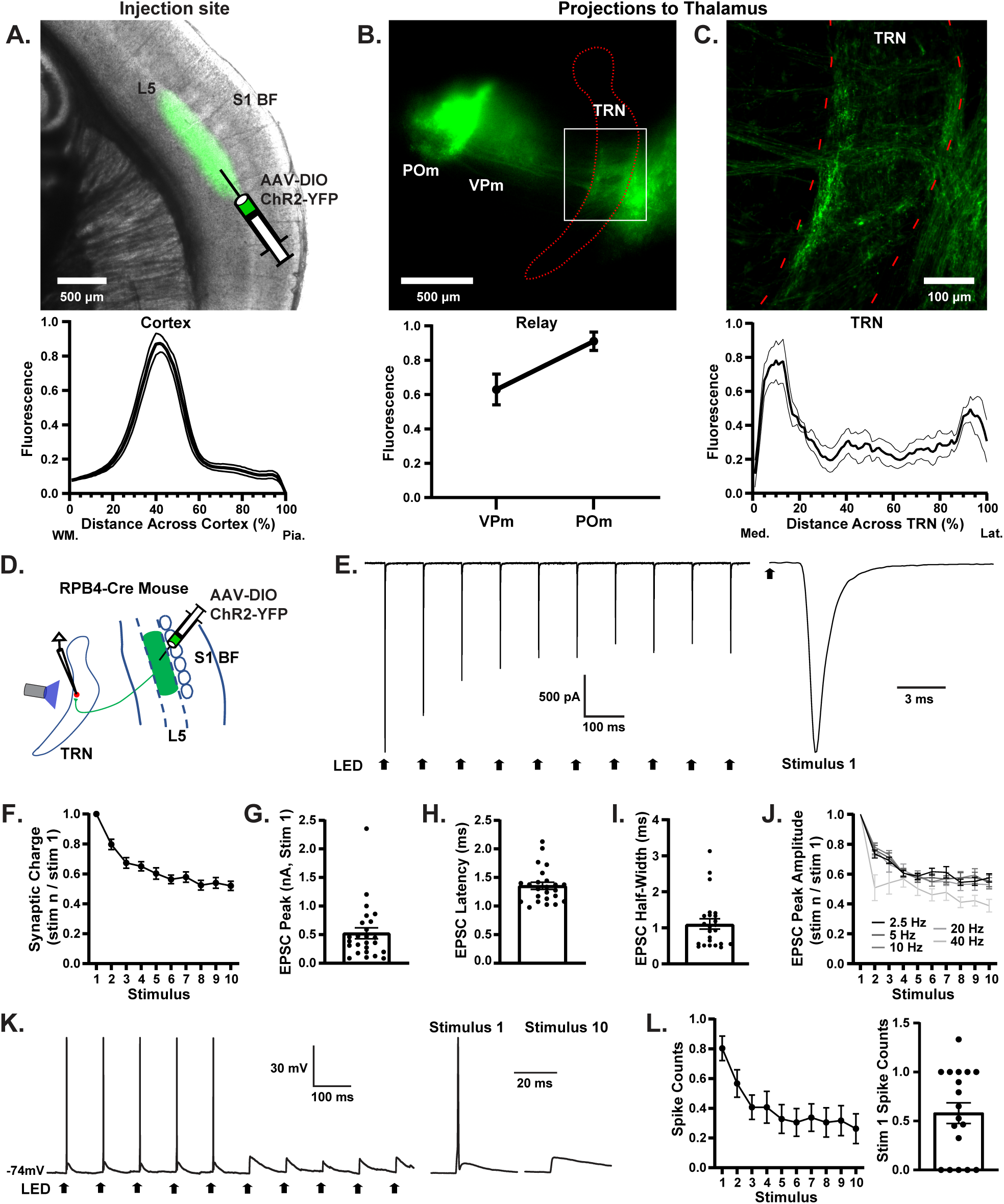
Monosynaptic input to HO TRN from L5 CT cells of S1. (A) **Top**, image of a live slice from an RBP4-Cre mouse in which Cre-dependent AAV1-DIO-ChR2-eYFP was injected into S1 barrel cortex to induce ChR2-eYFP expression in L5 cells (eYFP shown as green). **Bottom**, group fluorescent profile across the cortical depth, peaking in L5 (mean ± SEM; n=12 mice). (B) **Top**, image from the same preparation as (A) illustrating the L5 CT projections in the thalamus (green). **Bottom**, mean normalized fluorescence of L5 CT axons in the thalamocortical nuclei. L5 projections were more concentrated in POm than VPm, consistent with known anatomy^71,81,83,125^ (mean ± SEM, n=12). (C) **Top**, magnified image of a 50 um thick section taken from the slice in (B) following immunohistochemistry for parvalbumin to outline the TRN (red dotted line) and YFP to identify L5 axonal projections (green). Dense L5 CT varicosities are seen near the medial and lateral edges of TRN. **Bottom**, group fluorescence profile of CT projections across the TRN (mean ± SEM, n= 7). (D) Diagram of experimental setup. L5 CT axons were optogenetically activated and postsynaptic responses were recorded in TRN cells near the fluorescing L5 axons in the medial edge zone. Optical stimuli were 10 Hz trains, 10 pulses/train, 0.1 ms pulse durations, 23.5-25.7 mW LED intensities. (E) **Left**, Representative EPSCs from a TRN neuron, evoked by train stimulation of L5 CT axons. **Right**, EPSC evoked by the 1^st^ stimulus of the train, with magnified time base. (F) Group plot of synaptic charge evoked by the L5 CT train stimuli (values are means ± SEM, normalized to the response to stimulus 1, n=24 cells from 12 mice here and for panels G-I). The mean charge of the EPSCs for stimulus 1 was 0.804 ± 0.105 pC (calculated over the initial 10 ms following offset of each stimulus). Note the synaptic depression, contrasting with the facilitation observed with L6 CT stimulation (see Figure 2). (G) Peak amplitudes of the EPSCs evoked by the 1^st^ stimulus in the train. Each point represents 1 cell. (H) EPSC onset latencies (from onset of stimulus 1). Latencies for all 24 cells were < 2.5 ms, consistent with monosynaptic responses. (I) Widths at half maximum of L5-evoked EPSCs. (J) Normalized EPSC amplitudes over 10-pulse LED trains for a range of stimulus frequencies. There is little difference in short-term synaptic depression between frequencies, except for 40Hz (n=11 cells). (K) Example traces illustrating depressing action potential pattern evoked by L5 CT input in an edge TRN cell, consistent with the synaptic currents in (E-J). Responses to the 1^st^ and 10^th^ stimuli are shown at magnified time base on the right. (L) **Left**, group plot of spike counts evoked per stimulus (mean ± SEM, n = 13 cells). **Right**, mean spike counts evoked by the 1^st^ stimulus of the trains for each cell. Of the 18 cells recorded in current clamp, spiking could be evoked in 13. Steady-state Vm for K-L was ∼-74 mV.

To determine whether these L5 projections make functional synapses in TRN, we stimulated them optically while recording from TRN neurons (Figure 4D). Indeed, clear synaptic currents were evoked in HO cells located along the medial edge of TRN, near the dense fluorescent terminal labeling (Figure 4C-J). Unlike EPSCs evoked by L6 CT synapses (which facilitated – Figure 2), the EPSCs evoked by L5 inputs to TRN typically depressed during repetitive activation (Figure 4E, F, J), similar to L5 CT-evoked responses in HO thalamocortical cells (Figure S6)^71,73,121–123,129^ and anterior TRN^68^. These EPSCs in TRN had monosynaptic characteristics, including fast onset latencies and short durations (Figure 4E, H-I). Also note that the thalamocortical nuclei were removed from these slices, preventing possible cortico→thalamo→reticular transmission, thus providing further evidence for the monosynaptic nature of the EPSCs recorded in TRN.

The S1 L5 inputs to HO TRN cells were robust, often capable of evoking postsynaptic action potentials (n= 13/18 cells in which it was tested), but generally for only the first few stimuli in a 10 Hz train (Figure 4K-L), consistent with the strong short-term depression of the synaptic conductances. Even for those cells in which maximal responses were subthreshold for action potentials, the average 1st EPSP amplitudes were 4.8 +/− 1.18 mV, and therefore capable of modulating excitability.

The robust monosynaptic activation of HO TRN cells suggests that L5 CT activity could lead to disynaptic inhibition in the targets of HO TRN. To test this possibility, we recorded cells in one of those targets, the POm, while stimulating L5 CT axons (Figure S6A). L5 CT activation often triggered relatively short latency inhibition in POm (Figure S6B-C, 6.65ms +/− 0.47ms SEM), consistent with a disynaptic mechanism mediated through HO TRN cells. Additionally, L5 CT activation evoked only sparse POm cell firing (Figure S6; n=6/16 cells), and the POm spiking that did occur had fairly long latencies (4.32ms +/− 1.01 SEM; Figure S6E). This argues that the observed inhibition occurs through a disynaptic mechanism (e.g., L5→TRN→POm), rather than a trisynaptic one (L5→POm→TRN→ POm).

### M1 CT projections to HO TRN

The motor cortex (M1) makes top-down projections to the TRN and multiple thalamocortical nuclei, including the POm^67,68,75,79,130–133^. These projections originate from both L5 (pyramidal tract) and conventional L6 CT neurons of M1.

CT systems generally target groups of TRN and thalamocortical neurons that are themselves interconnected (e.g., L6 CT cells of V1 innervate the visual sector of TRN and the dorsal lateral geniculate)^11,12,14,19^. This raises the possibility that M1 may target HO neurons of somatosensory TRN, as these TRN neurons are known to bidirectionally synapse with the POm^11,27,60^. Such an arrangement would provide M1 with a degree of feedforward inhibitory control over POm.

Given the M1 CT results reviewed above, including reports from recent L5 CT studies^67,68^, it seemed possible that M1 CT systems of L5 or L6 could provide synaptic input to somatosensory TRN. Thus, we chose to cast a wide net, targeting both types of M1 CT pathways by stereotaxically injecting AAV2 carrying ChR2 genes in a recombinase-*independent* fashion across layers 5-6 of M1. We then tested for synaptic effects in TRN (Figure 5). Cortical ChR2-eYFP expression was centered on the vibrissa region of M1 (vM1)^97,134–136^ (Figure 5A). The projections to the somatosensory thalamocortical nuclei were most dense in POm (Figure 5B), as expected from previous reports (above). Projections from M1 to the TRN were centered on anterior regions of the nucleus, but often reached as far posterior as the somatosensory sector. On average, the projections to TRN reached –1.29 mm posterior to bregma (range –1.1 to –1.4 mm, n=7). Note that the somatosensory sector is located from approximately –1.1 to –1.7 mm posterior to bregma^60^. Within the somatosensory sector of the TRN, the highest projection densities were generally located in the medial HO zone (mean peak fluorescence: 17.1% of the distance from the medial to the lateral edge of TRN; Figure 5B).

**Figure 5.**
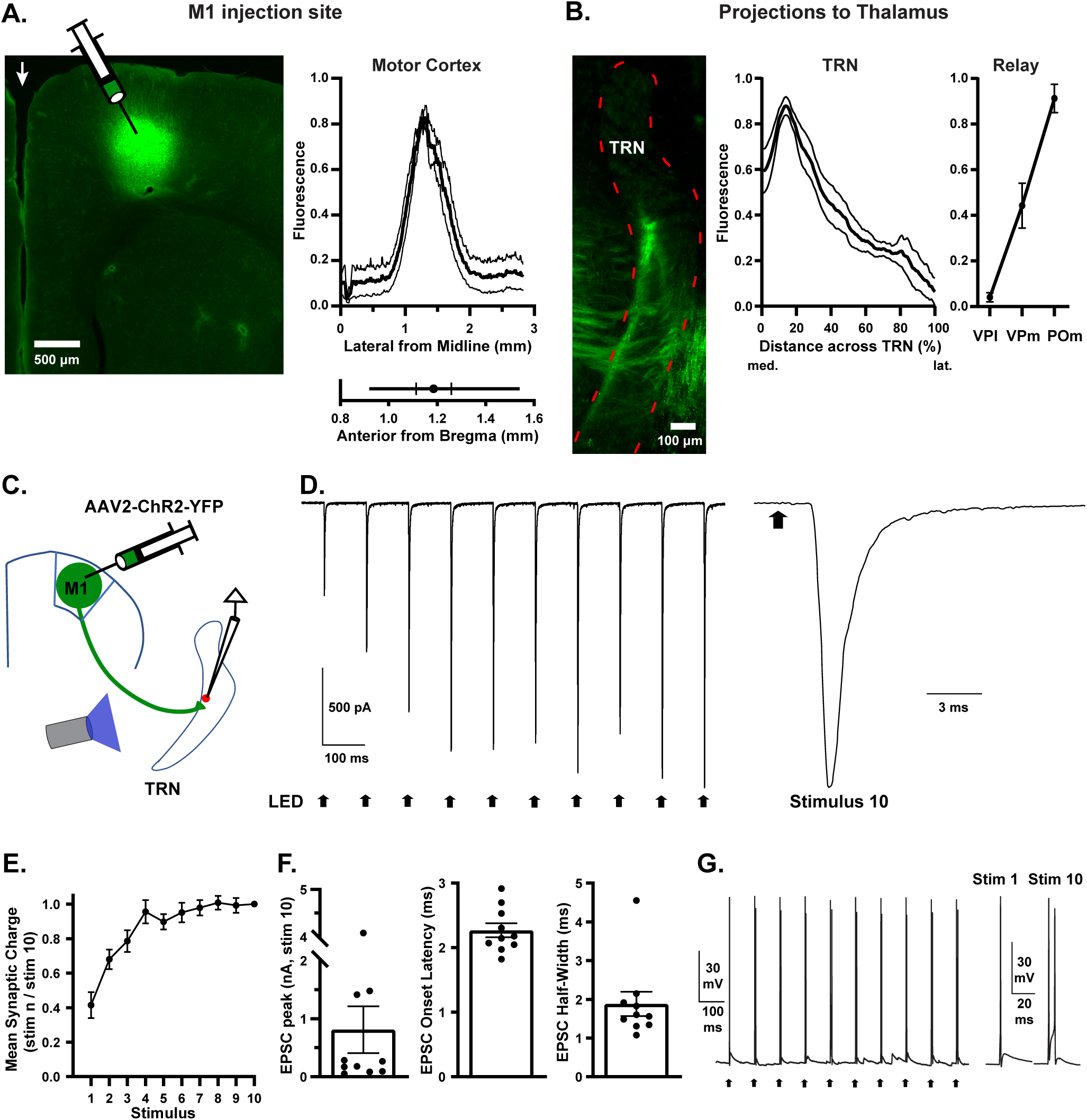
M1 CT synaptic input to HO somatosensory TRN. (A) **Left**, image of a coronal slice though frontal cortex from a mouse in which AAV2-ChR2-eYFP was injected into infragranular layers of vibrissa motor cortex to induce ChR2-eYFP expression in CT cells. Peak cortical eYFP expression in the example is approximately 1.3 mm lateral and anterior to Bregma. Arrow at midline. **Right**, mean group (± SEM, n = 7 mice) cortical fluorescence along the M-L axis (top), and the mean A-P location of peak labeling (bottom; hash marks = SEM, line length = range). The location of peak labeling (mean ± SEM, in mm) was 1.37 ± 0.06 lateral and 1.18 ± 0.07 anterior to bregma, at a depth of 0.68 ± 0.08 from the pial surface (53.4 ± 6.4% of the distance from the pia to w.m.). (B) **Left**, example of M1 projections in TRN (green). TRN was outlined (red dashed line) using immunohistochemical staining for parvalbumin. **Middle**, mean (± SEM; n=10) fluorescence profile of M1 projections across TRN, peaking in the HO medial edge zone. **Right**, Relative densities of M1 projections to the somatosensory thalamocortical nuclei. Projections were more concentrated in POm than in VPm or VPl, consistent with previous observations (e.g., ^66,81,132^). (C) Diagram of experimental setup. ChR2 was induced in M1 CT cells with recombinase-independent AAV2-ChR2-eYFP and CT axons were optically activated. Postsynaptic responses were recorded in TRN cells near the fluorescing M1 terminals in the medial edge zone. Optical stimuli were 10 Hz trains, 10 pulses/train, 0.1 ms pulse durations, 15-24 mW LED intensities. (D) **Left**, EPSCs from an HO edge TRN cell evoked by M1 CT stimulation. **Right**, EPSC evoked by the 10^th^ stimulus in the train, with magnified time base. (E) Group plot of synaptic charge evoked by the M1 CT train stimuli (means ± SEM, normalized to the response to stimulus 10; n=10 cells from 7 mice, here and for panel F). The mean postsynaptic charge for stimulus 10 was 1.543 ± 0.740 pC (calculated over the initial 10 ms following stimulus offset). Note the synaptic facilitation, consistent layer L6 CT synapses (see Figure 2 and related text). (F) Kinetics of EPSCs for the 10^th^ stimuli in the trains, measured for the same 10 cells as in E. **Left**, EPSC amplitudes (each point represents 1 cell). **Middle**, EPSC latencies. All were < 2.6 ms, consistent with monosynaptic responses. **Right**, EPSC widths at half maximum. (G) **Left**, example traces of action potentials evoked by M1 stimulation in an edge TRN cell. **Right**, responses to 1^st^ and 10^th^ stimuli, with magnified time base. The facilitating spiking patterns are consistent with the synaptic currents in (D-E). Steady-state Vm was approximately –74 mV.

Optogenetic activation of the M1 CT terminals evoked clear monosynaptic EPSCs in TRN cells located near the dense fluorescent terminal labeling in and around the medial HO zone (11/11 cells; Figure 5C-F). On average, recorded cells were located 15.7% of the distance from the medial to lateral boundary of TRN (range 2.5-25.5%) and 1.15 mm posterior to bregma (range 1.40 to 0.95 mm posterior), which is slightly anterior to the typical center of CT projections from S1 barrel cortex (by ∼0.25 mm)^60^, but well within convergent range (see below).

Synaptic responses evoked by activation of M1 CT synapses varied broadly in amplitude (mean EPSCs = 0.81 nA; range 0.054 – 4.082 nA when stimulated at 15.54mW LED intensities; Figure 5D-F), and were sometimes strong enough to trigger action potentials (2/5 cells tested in current clamp mode; Figure 5G). EPSC latencies and durations were short, consistent with monosynaptic connections (Figure 5D, F), and these responses facilitated with repetitive activation in 10/11 cases, consistent with L6 CT synapses (Figure 5D-E).

### Neurons in HO TRN integrate convergent synaptic input from S1 L5 and L6 CT systems

The projection patterns from S1 L5 to TRN were largely complementary to those from S1 L6, but there appeared to be overlap in the HO zone near the medial edge of TRN (Figures 1A-B; 4A-C; S1B-E). This raised the possibility that the L5 and L6 CT inputs may converge on individual TRN neurons located in this overlap zone, allowing for interactions in the influences of these distinct CT pathways.

To explore this possibility, we made dual AAV injections in Ntsr1-Cre mice to express ChrimsonR-GFP in the L6 CT pathway (using a Cre-dependent AAV construct) and ChR2-mCherry in the L5 CT pathway (using an FAS “Cre-Off” construct in which ChR2 expression depended on the absence of Cre recombinase)^137^(Figures 6A, S7A).

**Figure 6.**
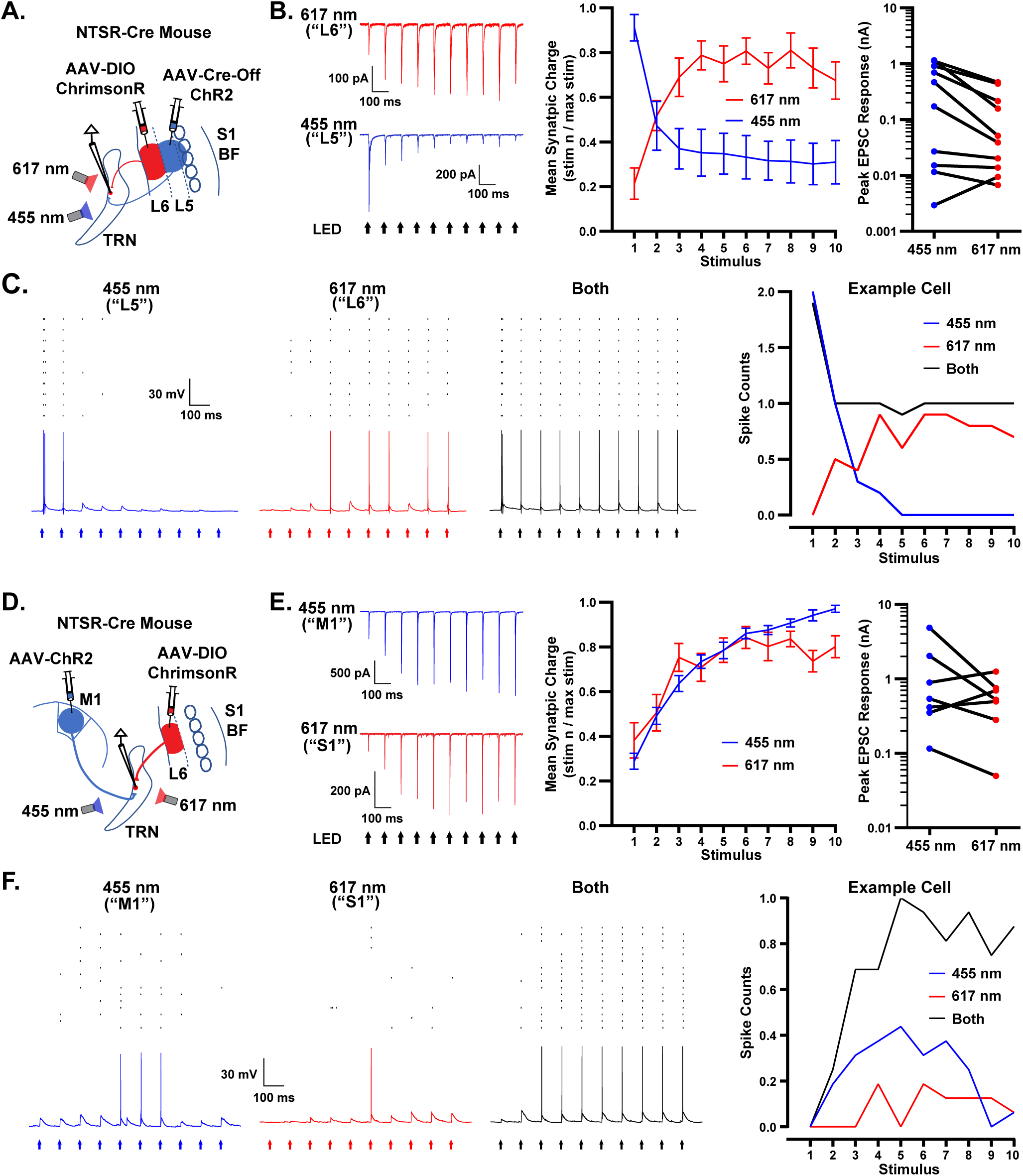
HO TRN neurons integrate convergent input from multiple CT pathways. (A) Diagram of experimental setup for tests of S1 L5/L6 CT convergence. Ntsr1-Cre mice were injected in the infragranular layers of S1 with two constructs. The first was a standard AAV-DIO, driving expression of ChrimsonR in Cre-expressing L6 CT cells (red). The second was an AAV Cre-Off construct, driving ChR2 expression in all neurons except those with Cre-recombinase; thus, ChR2 was expressed in L5 CT cells (blue) but excluded from L6 CT cells (because they were Cre-positive). Recordings were performed in TRN where the L5 and L6 CT projections converged (the HO medial edge zone) while alternately stimulating with a 455nm LED (activating ChR2-expressing L5 axons) and a 617nm LED (activating ChrimsonR-expressing L6 axons). (B) **Left**, example of convergent L6 and L5 CT input to a TRN cell; EPSCs evoked by alternating 617 nm and 455 nm stimuli (Vhold –84mV). **Middle**, mean synaptic charge evoked by each stimulus, normalized to the largest responses in the trains (± SEM; n=10 cells from 5 mice; charge integrated over the 10 ms following the stimulus). **Right**, peak EPSC amplitudes evoked by 455nm and 617nm stimuli. Lines connect responses for the two classes of CT input for individual TRN cells (same n=10 cells from 5 mice as middle panel). (C) **Left**, example action potential (AP) responses evoked by L5 input (455 nm), L6 input (617 nm), or both inputs simultaneously (steady-state Vm –74mV). Raster plots above the traces show AP times for each sweep. **Right**, mean spike counts from the recording shown on the left. Note the depression to L5 input, the facilitation to L6 input, and the more dynamically stable summated response. (D) Diagram of experimental setup for tests of M1/S1 CT convergence. AAVs were injected into infragranular layers of M1 and S1 to drive expression of ChR2 in M1 CT cells (blue) and ChrimsonR in S1 CT cells (red). The AAV constructs in M1 were recombinase independent (AAV2-ChR2-EYFP; 4 mice), as were those in S1 for 2 of these mice (AAV2-ChrimsonR-tdT). For the other 2 mice, the AAVs in S1 were Cre-dependent (AAV2-DIO-ChrimsonR-tdT, NTSR1-Cre genotype). Whole-cell recordings were performed in TRN where the M1 and S1 CT projections converged (the HO medial edge zone), while alternately stimulating with a 455nm LED (activating ChR2-expressing M1 axons) and a 617nm LED (activating ChrimsonR-expressing S1 axons). (E) As for panel B except illustrating convergent M1 (blue) and S1 (red) CT input to TRN (n=7 cells from 4 mice). The EPSCs evoked by both M1 and S1 inputs underwent facilitation. (F) As for panel C except responses are evoked by M1, S1, and combined M1/S1 inputs. Note the summation and facilitation of spiking with the combined stimulation.

In these dually-injected mice, ChrimsonR-GFP was expressed in the L6 cells of S1 and their projections to the thalamus/TRN, consistent with the previous Cre-dependent expression in Ntsr1-Cre mice (compare ChrimsonR-GFP expression in Figure S7A-D with ChR2-eYFP expression in Figure 1A-B). As expected, there were clear projections to primary central TRN and the HO zone near the medial edge (Figure S7B-C, green). ChR2-mCherry was more broadly expressed in S1 across laminae; notably, there was fairly strong labeling in layer 5 and much weaker labeling in L6, consistent with exclusion from the Cre-expressing NTSR1 cells in the latter (Figure S7A, red). The ChR2-mCherry expressing projections to thalamus in these mice resembled the L5 CT projections observed in RBP4 mice, including relatively strong input to the HO edge zones of TRN, and much weaker input to the primary central zone (compare Figure 4C with S7B-C). Importantly, there appeared to be overlap of fluorescently labeled terminal zones of the L5 and L6 CT pathways near the medial edge of TRN (Figure S7B-C).

Neurons located at the intersecting terminal zones of the two CT pathways in TRN were tested physiologically for convergent synaptic input. A 617 nm LED was used to activate ChrimsonR-expressing NTSR/L6 CT axons and a 455 nm LED was used to activate ChR2-expressing NTSR-negative L5 CT axons (Figure 6A). The opsin specificity of the responses was verified using a control procedure^138^, indicating that the spectrally separate LEDs indeed activated largely distinct ChrimsonR– and ChR2-expressing axonal populations (Figure S8). Among the TRN cells tested for dual S1 L5 and L6 innervation, 10/10 were found to receive convergent input (Figure 6A-B). The synaptic responses evoked by the 617 nm LED facilitated during repetitive stimulation, as expected for a L6 CT pathway (10/10 cells). In the same cells, responses evoked by the 455 nm LED generally depressed with repetitive stimulation, consistent with a L5 CT pathway (9/10 cells)(Figure 6B).

When TRN cells were tested in current clamp mode, integration of convergent L5 and L6 inputs led to stronger spiking responses, with somewhat more “stationary” patterns during repetitive stimulation, than either class of input alone (Figure 6C). The latter is presumably due to the net/compound synaptic conductances being more stable than those of the component pathways, which individually undergo strong short-term depression (L5) or facilitation (L6), respectively (Figure 6B and Figures 4, 2). More generally, these results indicate that the distinct CT pathways can act in concert to influence the activity (and consequently the inhibitory output) of neurons in the higher order region of somatosensory TRN.

### M1 and S1 L6 CT inputs also converge on and summate in HO TRN neurons

Like L5 of S1, the projections from M1 appeared to overlap with those from S1 L6 in the medial HO zone of TRN (Figures 1A-B; 5A-B; S1B-E). Again, this raised the possibility of functional integration in TRN neurons within the overlapping zone.

To test for this, we made separate AAV injections in infragranular layers of M1 and S1. AAVs carrying recombinase-independent ChR2-EYFP genes were targeted to the infragranular layers of M1 as described previously (Figure 5). During the same surgery, AAVs carrying either recombinase-independent ChrimsonR-tdT or Cre-dependent ChrimsonR-tdT genes were targeted to L6 of S1; the Cre-dependent AAVs in S1 were applied in Ntsr1-Cre mice (see Methods). We then tested for synaptic convergence and integration of the M1 and S1 CT projections to TRN. The expression patterns and projections to the thalamus in the dual S1 and M1 experiments were generally consistent with results from the single cortical injections described above (compare Figures 1A-B, S1B-E and 5A-B with Figure S7E-I). As expected, there was overlap of labeled terminals of the M1 and S1 CT pathways in the medial region of TRN (Figure S7G-H).

All TRN cells recorded from the region with anatomical convergence from M1 and S1 responded to both classes of CT input (8/8; Figure 6E). As expected from analysis of the individual pathways described previously (Figures 2, 5), TRN cell responses evoked by both M1 (8/8) and S1 L6 (7/8) underwent short-term facilitation, consistent with L6 CT synapses (Figure 6E). As with the L5/6 inputs, recordings of TRN cell responses to simultaneously activated S1 L6 and M1 pathways revealed functional integration, with clear enhancement of TRN spiking to combined M1/S1 input compared with either pathway alone (Figure 6F).

## Discussion

This study reveals diverse influences of the neocortex on primary and HO neurons of the TRN, with implications for top-down, state-dependent, regulation of processing in sensory thalamocortical circuits. We demonstrate that the primary and HO sensory TRN subtypes receive unique patterns of cortical inputs that exert varied effects on their activity due to specializations in both synaptic and cell-intrinsic properties.

### Patterns of CT inputs from L6 of S1 to primary and HO zones of somatosensory TRN

Top-down communication between the neocortex and the TRN has traditionally been thought to occur exclusively via L6 CT projections from functionally aligned cortical areas^2,10–12,16,19,50,65^. Therefore, our investigations of neocortical influences on the somatosensory TRN initially focused on CT inputs from L6 of S1.

Anatomical experiments, using Cre-dependent AAV constructs in NTSR1-Cre mice, revealed dense S1 L6 innervation of the primary central TRN, with less intense projections to the HO medial edge TRN. Using a within-mouse dual tracing approach, we further demonstrated that the projections to primary and HO TRN originated from CT neurons located in distinct sublaminae (upper and lower L6, respectively), consistent with conclusions from previous studies^80,83^.

The barrel subfield of S1 processes sensory inputs mainly from the head, whereas non-barrel subfields (e.g., forelimb, hindlimb, trunk) handle inputs from other parts of the body. Here we observed that the L6 CT pathways from barrel and non-barrel S1 target distinct regions of primary TRN, with dense but spatially separate projections^83,91^. This suggests that primary TRN cells receive their main cortical input from either barrel or non-barrel CT pathways, but not both. In contrast, the barrel and non-barrel L6 projections to HO TRN were less dense and overlapped spatially, suggesting convergence on HO TRN cells. These contrasting motifs – highly specific and focused CT influences on primary TRN cells versus more widespread and distributed influences on HO TRN – are intriguing and appear to extend even beyond the canonical S1 L6 CT systems, as discussed below.

Previous work has shown that the POm thalamus provides robust synaptic input to HO TRN neurons in both the medial and lateral edge zones of somatosensory TRN^60^. In the present study, CT projections to HO TRN were clearly more intense to the medial than the lateral zone^80^. However, we previously recorded clear EPSCs in lateral HO TRN in response to S1 CT input^139^; in that study, the laminar identity of the CT projection was unspecified. It will be important to fully characterize the origins and effects of CT inputs to the two groups of HO TRN neurons in future studies.

### Integration and functional impact of L6 CT projections from S1

Our optogenetic-electrophysiological experiments revealed that synaptic currents evoked by S1 L6 CT inputs were more than two-fold larger in primary than HO TRN cells, consistent with the stronger anatomical projections. These greater currents contributed to far stronger CT-evoked spiking in primary cells. However, differences in input strengths did not fully account for the variations in spike rates among TRN cell subtypes. An even more potent factor was a property intrinsic to the TRN cells. Specifically, primary TRN cells had much stronger T-type calcium currents^23,27,59,60,62^; also see ^58,98–101^, which were found to summate with synaptic currents and boost their spike responses to L6 CT inputs.

The T-type currents not only affected response magnitudes, but they also shaped dynamic patterns of spiking by triggering bursts during initial phases of responses, followed by gradual decreases in bursting, likely due to voltage-dependent inactivation^59,60,89,102–104,106^. In the primary TRN cells, this resulted in strong phasic L6 CT-evoked spiking that depressed during repetitive stimulation, overcoming the facilitating nature of the synaptic inputs. This phasic pattern of primary TRN spiking aligns with a role for these cells in rapid, modality-specific sensory processing. This pattern can be critical for the frequency-dependent balance of excitatory and inhibitory influences induced by L6 CT inputs in primary TC relay cells^8,89,140,141^ (also see ^90,142,143^). The absence of such a pattern in HO TRN cells is notable, and suggests that CT feedforward inhibition in the distinct thalamic subcircuits may be tailored to specific functional roles.

### State dependence

Thalamic circuits are under strong regulatory influences of arousal-related brain states^113,115,116,119,144–153^ – states which affect behavioral/perceptual performance^118^. Neurons of the TRN participate in this regulation; their activity is affected by behavioral arousal and attentional demands^16,20,22,26,32,33,57,154,155^, and their synaptic outputs to thalamic neurons likely play important roles in many processes associated with those states^23,25,29–31,43,57,156–158^.

Classically, neurons in TRN have been observed to fire in high-frequency bursts (>300 Hz) during quiescent states, and in lower frequency “tonic” patterns during arousal^11,110,114,154,159–161^. These firing modes are thought to depend on the properties of the T-type calcium channels investigated above^102–104,106^. During quiescence, thalamic cells have hyperpolarized resting potentials, allowing the T-type channels to deinactivate. Under those conditions, phasic stimuli can trigger large T-type calcium currents which drive spike bursts, like the bursts in primary TRN evoked by CT stimuli. Transitions to aroused states are associated with increased release of modulatory transmitters that cause depolarization, inactivation of T-type channels, and tonic firing^18,35,98,111,112,117,155,162–165^.

Here we found that shifts in steady-state voltage, which mimic those occurring during arousal, dramatically alter S1 L6 CT-driven firing patterns in primary TRN. However, the same voltage manipulations had little effect on firing in HO TRN cells; the latter expressed facilitating CT-evoked responses (with little bursting) no matter the steady-state potential, consistent with a lack of strong T-type calcium currents.

The differences in bursting may have important consequences for state-dependent CT inhibition mediated by the two TRN systems, including on magnitudes and spatiotemporal patterns of inhibition^14,89,166–168^. CT-regulated TRN bursting is pivotal for pacing spindle oscillations, a prominent rhythm of non-REM sleep^11,14,30,59,169–173^, which is often disrupted in neurological disease and may be important for learning-related plasticity^34–37,39,40,48,116,174–176^. A number of reports have identified specific TRN cell types or anatomical regions as preferentially involved in spindles, with a common feature among them being a strongly bursting phenotype^23,27,31,59^.

But what are the roles of non-bursty TRN neurons, such as the HO cells described here^27,59,60^? Intriguingly, Halassa and colleagues reported that activity in the limbic sectors of TRN (which others have found to have little bursting^23^), was suppressed during sleep/spindles but elevated during arousal – opposite to sensory TRN which was most active during sleep/spindles^57^. The physiological similarities between limbic TRN cells and HO cells of the somatosensory TRN studied here raise the possibility that they may have similar functions in state-dependent network activity and top-down CT feedforward inhibition of thalamocortical signaling.

### Convergence of multiple CT pathways on HO TRN neurons

In addition to receiving modest convergent inputs from L6 of both barrel and non-barrel cortices, our findings demonstrate that HO TRN neurons also integrate inputs from other CT pathways, including those originating from L5 of S1 and from M1. This indicates that HO TRN is involved in processing diverse types of top-down sensory and motor signals, potentially facilitating coordination among the distinct CT systems.

Traditionally, L5 CT projections have been thought to bypass the TRN; however, a select group of studies have challenged this view, reporting that L5 CT cells from *frontal areas* make synapses in anterior sectors of the TRN^66,68^ (also see ^32,69^). Our findings, together with an anatomical study from Sherman and colleagues^67^, expand on this new insight, demonstrating that L5 neurons from *sensory areas* also form synapses with TRN, particularly with cells in its HO zones.

Our optogenetic-physiological experiments verified and characterized the functionality of these L5 CT synapses. EPSCs evoked by layer 5 inputs were fast and depressed during repetitive activation – hallmarks of layer 5 CT-evoked responses recorded in other systems and cell types^68,71,73,121–123,129^. These kinetics suggest a mechanism for rapid dynamic control of HO TRN by the neocortex. In fact, L5 inputs were often robust enough to drive action potentials in HO TRN cells, and seemed to trigger feedforward inhibition in POm.

Our dual pathway experiments further revealed that individual cells of HO TRN receive convergent input from both L5 and L6 systems. Summation of these distinct inputs not only increased overall spike rates, but also affected the dynamics of HO TRN activity. During repetitive stimulation, L5 inputs alone evoked phasic/depressing responses, while L6 inputs alone evoked facilitating responses. However, simultaneous activation of both pathways resulted in more stable spike patterns. This was likely caused by the relative stability of the summated conductances evoked by the combined inputs during repetitive stimulation.

It is worth noting that our observations of sensory L5 projections to TRN contrast with findings from a recent study by a prominent research group which reported that L5 CT projections to TRN are essentially limited to frontal systems^68^. Resolving this issue will be an important focus for future research.

Similar to L5 of S1, the projections from M1 also targeted the HO zone along the medial edge of somatosensory TRN. The methods applied in our M1 experiments did not isolate projections from specific cortical laminae. However, the recorded synaptic currents exhibited facilitation during repetitive activation, consistent with L6 rather than L5 CT synapses. It is possible that L5 cells of M1 may also form synapses in somatosensory TRN, as they do in anterior TRN^67,68^. This can be resolved in future studies using methods that isolate L5 projections from M1.

Nevertheless, it is clear that M1 and S1 CT projections converge on HO TRN cells, where their inputs summate. This convergence was observed in approximately the same medial HO region of TRN as the L5 and L6 projections from S1. Based on this it seems reasonable to speculate that a population of HO TRN neurons integrate input from all three CT systems. The functions of this convergence are yet to be determined, but it provides a mechanism by which the distinct CT pathways from various domains could act in concert to regulate the activity and inhibitory influences of HO TRN cells, potentially supporting processing of complex sensory information essential for higher order functions.

Growing evidence suggests that excitatory neurons in HO TC nuclei can serve as “transthalamic relays” for communication from primary to HO sensory cortical areas^51,54,77,151,177,178^. In this model, L5 CT cells in primary areas send processed cortical information to HO TC cells, which then relay it to HO cortical areas, including motor cortex^179^. Similarly, motor cortex might send copies of efferent motor commands to sensory areas transthalamically, in addition more direct routes. CT feedforward inhibition could be an ideal mechanism to gate the activity of TC neurons carrying information between cortical areas.

Moreover, the high degree of convergence from different CT systems to HO TRN cells provides a possible mechanism for crossed inhibitory gating in which one cortical system may be able to regulate signaling from another^12,14–17,180^.

It is striking how the convergent CT inputs to HO TRN mirror the known inputs to the HO TC neurons that these TRN cells inhibit. For instance, the same CT areas and layers found to project to HO somatosensory TRN also project to one of its main targets, the POm (reviewed above and in ^5,8,10,75,81^). This raises the possibility that the excitatory effects of CT projections to HO TC cells might be balanced by feedforward inhibition through HO TRN. It will be important to understand how this feedforward inhibition interacts with inhibition mediated by extrathalamic sources, such as the zona incerta and anterior pretectal nucleus, which are also innervated by L5 CT cells^14,71,181^.

### Future Directions

The influences of the TRN cell subtypes in top-down CT processes are ultimately mediated via disynaptic feedforward inhibition at their targets in the primary and HO TC nuclei^8,14^. The characteristics of that inhibition, and the ways it may vary between subcircuits, will depend not only on how activities of the TRN subtypes are controlled by their CT afferents (as in the present study), but also on the properties of their GABAergic synaptic outputs. Therefore, it will be important to characterize the fundamental features of the synapses, such as their morphologies, transmitter release probabilities and dynamics, and features of the GABA receptor subtypes. These factors can significantly impact the strengths, kinetics, and dynamic patterns of inhibition in TC cells, including feedforward inhibition triggered by CT inputs^14,35,89,166–168,182–186^. In multiple systems, the balance of excitation and inhibition (E/I balance) evoked by CT inputs has been shown to depend on timing and/or frequency of corticothalamic activity^89,90,140–142^, the effects of which seem to be mediated, in part, by short-term plasticity of the GABAergic synapses linking the TRN and TC cells^89^. Clarification of the basic features of these inhibitory synapses will undoubtedly facilitate our understanding of the mechanisms of this temporally-sensitive E/I balance and other functional specializations of CT feedforward inhibition mediated through primary or HO TRN.

*In vitro* methods are appropriate for probing the fundamental organization, as well as cellular and synaptic mechanisms, of the CT → TRN pathways under investigation. To broaden our understanding of CT feedforward inhibition at a functional level, *in vivo* experiments in awake mice will be critical. Intense and sharply timed optogenetic stimulation as used here has the potential to induce levels of CT synchrony exceeding that during natural activity. Indeed, natural L6 CT firing is generally sparse, but activity can be dynamic and heterogenous in certain sensory or behavioral conditions^145,148,187–190^. To investigate CT actions under conditions less prone to synchronization, alternative strategies can be employed to manipulate and infer the effects of top-down CT projections. This could include application of lower intensity sustained optogenetic modulation with the goal of altering excitability of CT cells while allowing normal afferents to drive the precise timing of CT activity (e.g., ^191^). Alternatively, other methods using multiphoton spatial light modulation techniques could permit unsynchronized activation of many individual CT neurons in concert^192,193^, with temporally precise patterns based on recordings of natural activity.

### Conclusions

The pathways interconnecting the neocortex, the TRN, and TC nuclei, have long been recognized as crucial for sleep, consciousness, and the perception of external signals. The emergence of the primary and HO framework for thalamocortical organization, which divides these pathways into distinct subsystems, has provided a valuable catalyst to refine our understanding. However, specific knowledge about the TRN neurons within these subsystems is only beginning to emerge. Our results uncover fundamental organizational and operational principles about cortical influences on primary and HO TRN, addressing an important gap in our understanding of thalamocortical functioning.

## Methods

All procedures were done in accordance with the NIH Guidelines for the Care and Use of Laboratory Animals and approved by the University of Alabama at Birmingham Institutional Animal Care and Use Committee.

### Animals

The following mouse lines were used: NTSR1-Cre (MMRRC Tg(Ntsr1-cre)GN220Gsat/Mmucd), RBP4-Cre (MMRRC Tg(Rbp4-cre)KL100Gsat/Mmucd), PV-Cre (The Jackson Laboratory, Pvalb-IRES-Cre, #008069), PV-Flp (The Jackson Laboratory, Pvalb-T2A-FlpO-D, #022730), CB-Cre (The Jackson Laboratory, Calb1-IRES-Cre-D, #328532), Ai14 (The Jackson Laboratory, Ai14(RCL-tdT)-D, #007908), Ai65F (The Jackson Laboratory, Ai65F(RCF-tdT), #032864), Ai32 (The Jackson Laboratory, Ai32(RCL-ChR2(H134R)/EYFP), #024109), Syt7KO (The Jackson Laboratory, Synaptotagmin VII knockout, #004950), ICR (Charles River, CD-1[ICR], Strain Code 022).

The specific applications of the mouse lines are described in the Results and figure legends. Generally, NTSR1-Cre mice were used to selectively express opsin in L6 CT projection cells by combining with either Cre-dependent AAV constructs injected into the cortex or by breeding them with Cre-dependent ChR2-eYFP reporter mice (Ai32)^86–90^. RBP4-Cre mice were likewise used to express opsins in L5 CT cells^66,68,127,128^. PV-Cre and PV-Flp mice were crossed with recombinase-dependent tdTomato reporter strains (Ai14 or Ai65F, respectively) to drive expression of fluorescent tdTomato in neurons across the somatosensory sector of TRN^60,194^. In a subset of experiments, PV-Flp x Ai65F mice were subsequently crossed with NTSR1-Cre mice to create an “NTSR1-Cre x PV-Flp x Ai65F” strain, allowing specific opsin expression in L6 CT neurons and fluorescent identification of TRN neurons within individual mouse brains. Syt7KO mice were used to test the role of synaptotagmin 7 in short-term facilitation in the L6 cortico-reticular pathway. ICR and CD1-Cre mice were used as controls.

All 85 mice in this study had ICR backgrounds, and both sexes were used for experiments. Mice were maintained on a 12 h:12 h light/dark cycle, group-housed, and provided food and water ad libitum.

### Stereotactic virus injections

Injections of adeno-associated viruses (AAVs) were made in layers 5 and 6 of somatosensory and motor cortices to drive opsin-fluorescent protein expression in targeted corticothalamic pathways. Mice were deeply anesthetized with vaporized isoflurane (∼2%), placed in a stereotaxic frame, then injected with ∼0.02ml of local anesthetic (2% Lidocaine HCL) under the scalp. Approximately 10 minutes after lidocaine administration, an incision was made in the scalp, and craniotomies were performed over somatosensory and/or motor cortices with a 0.5mm drill bit. Glass micropipettes were used to pressure-eject viral solutions (0.1 ul to 2 ul) at a rate of ∼1ul/30min into cortical regions of interest in the right hemisphere. The pipette was held in its final position for 10-15 min after injection before withdrawal from the brain. The scalp was sutured with cyanoacrylate glue, subcutaneous carprofen (0.25mg/g) was administered for analgesia, then mice were recovered on a 37 ℃ heating pad for 1 hour before being returned to their home cages. Experiments were usually performed 9-12 days after the virus injections to allow for sufficient opsin/fluorescent protein expression.

Adeno-associated viruses (AAVs) were acquired from the University of North Carolina, Addgene or the University of Pennsylvania Vector Cores and used at the following titers, diluting from stock concentrations with sterile saline:

- AAV1-DIO-ChR2-eYFP: AAV1-EF1a-double floxed-hChR2(H134R)-EYFP-WPRE-HGHpA (titer = 2.8 to 9.9×10^12 vg/ml, Addgene).
- AAV2-DIO-ChR2-eYFP: AAV2.syn.diochr(134)eyfp.wp.hgh (titer = 4.5 to 6.0×10^12 vg/ml, Penn).
- AAV2-DIO-ChR2-mCherry: AAV2/Ef1a.DIO.hChRr2(H134R)-mCherry.WPRE.hGH (titer = 6.0×10^12 vg/ml, UNC).
- AAV2-ChR2-eYFP: rAAV2/hsyn-hChR2(H134R)-eYFP-WPREpA (titer = 3.7 to 5.6×10^12 vg/ml, UNC).
- AAV2-CreOff-ChR2-mCherry: AAV2.EF1a.FAS.hChR2(H134R)-mCherry-WPRE.hGH (titer = 5.7×10^12 vg/ml, Penn).
- AAV8-DIO-ChrimsonR-GFP: rAAV8-EF1a1.1-FLEX-ChrimsonR-GFP (titer = 1.9×10^12 vg/ml, UNC).
- AAV2-DIO-ChrimsonR-tdT: AAV2-Syn-FLEX-ChrimsonR-tdT (titer = 6.0×10^12 vg/ml, UNC).
- AAV2-ChrimsonR-tdT: AAV2/Syn-ChrimsonR-tdT (titer = 3.7×10^12 vg/ml, UNC).

To test projections from specific corticothalamic pathways to the TRN, viral solutions were injected into infragranular layers of selected cortical areas between postnatal day 11 and 18. Viral titers, expression times, and volumes were adjusted to achieve strong, on target expression while minimizing overexpression damage and nonspecific leakage.

To test synaptic inputs mediated by S1 barrel field (BF) L6 projections to the TRN (Figures 1-3, S1, S2), NTSR1-Cre (n = 8) or NTSR1-Cre x PV-Flp x Ai65 (n = 8) mice were injected with AAV1-DIO-ChR2-eYFP (above; 0.66 μl – 1.5ul volumes) at 3.4 mm lateral and 0.5 mm posterior from bregma, distributed evenly from 0.8 mm to 1.4 mm below the pia (x= 3.4mm, y= –0.5mm, z= 0.8mm to 1.4mm). For forelimb/hindlimb/trunk S1 L6 projections (Figure S1D-E), NTSR1-Cre x PV-Flp x Ai65 mice (n = 7) were injected with AAV1-DIO-ChR2-eYFP (1-1.5 ul) at x= 2.0mm, y= –0.65mm, z= 0.8mm to 1.2mm.

To test S1 BF L5 projections (Figure 4), RBP4-Cre mice (n=14) were injected with AAV1-DIO-ChR2-eYFP (1-1.5 ul) at x= 3.4 mm, y= –0.05 mm, z= 0.5mm to 1.2mm.

To test M1 projections to TRN (Figure 5), PV-Cre x Ai14 (n=4), NTSR1-Cre (n=2), or Calbindin-Cre mice (n=2) were injected with recombinase-independent virus (AAV2-ChR2-eYFP; 0.33 – 0.66 μl) at x= 1.25mm, y= 1.1mm, z= 1mm. In a subset of these M1 experiments, L6 of S1 BF was also injected with AAV carrying ChrimsonR-tdT genes, to allow testing of the convergence of M1 and S1 CT pathways onto HO TRN cells using separate wavelengths to activate distinct pathways (Figures 6, S7, S8D-E) (0.33 – 1 ul at x= 3.4mm, y= –0.5mm, z= 0.8mm to 1.4mm; AAV2-DIO-ChrimsonR-tdT used in the NTSR1-Cre mice, n=2; AAV2-ChrimsonR-tdT used in the Calbindin-Cre mice, n=2).

A similar dual opsin strategy was used to test for possible convergence of L5 and L6 CT pathways from the S1 BF to HO TRN (Figures 6, S7, S8A-B). In these experiments, Cre-Off and Cre-On constructs were used in NTSR1-Cre mice (n=5) to express ChR2 and ChrimsonR in L5 and L6 CT cells, respectively. In dual L5/L6 injections the stereotactic arm was adjusted so that the injector pipettes could be inserted perpendicular to the surface of the brain at the craniotomy (∼43° from vertical and revolved 10° clockwise), allowing for more accurate targeting of cortical layers. Craniotomies were made at x= 3.95 mm and y= –0.50mm from bregma, then L5 injection pipettes were inserted 0.7mm into the cortex, and 0.33 to 0.66 μl of Cre-Off AAV2-ChR2-mCherry was injected (at 5.7 x 10^12 vg/ml) to express ChR2 in L5 CT cells. Subsequently, L6 injection pipettes were inserted 1.1mm into the cortex, and 0.33 to 0.66 μl of AAV8-DIO-ChrimsonR-GFP was injected (at 1.9 x 10^12 vg/ml) to express ChrimsonR in L6 CT cells.

Finally, to compare the TRN projection patterns from upper and lower L6 CT cells of the S1 BF, dual L6 experiments (Figure S1) were conducted in NTSR1-Cre mice (n=18). Small volumes (0.33 μl) of distinct viral constructs were delivered to upper and lower L6 to isolate expression within the sublayers. Again, craniotomies were made at x= 3.95 mm and y= –0.5mm from bregma and injection pipettes were inserted perpendicular to the surface of the brain. Injection depths were ∼0.8 and 1.3mm from the cortical surface for upper and lower L6 injections. Viruses were counterbalanced so that AAV2-DIO-ChrimsonR-tdT was injected into upper L6 and AAV2-DIO-ChR2-eYFP into lower L6 in n=7 mice, while n=8 mice were injected in the opposite configuration. In n=3 mice (used exclusively for anatomy) AAV2-DIO-ChR2-YFP was injected into upper L6, and AAV2-ChR2-mCherry into lower L6. Titers were adjusted to achieve approximately equal cortical fluorescence.

Control experiments were conducted to test the specificity of activating ChR2-expressing and ChrimsonR-expressing terminal arbors with 455 nm and 617 nm LED stimuli (Figure S8)^138^. NTSR1-Cre mice were injected in L6 of S1 with either AAV1-DIO-ChR2-eYFP or AAV2-DIO-ChrimsonR-tdT (∼1 ul volumes; x= 3.4mm, y= –0.5mm, z= 0.8mm to 1.4mm).

### Slice preparation

After allowing time for opsin expression (typically 9-12 days), acute somatosensory thalamocortical brain slices were prepared for *in vitro* recording, as described previously^60,89^. These slices contain the somatosensory thalamus, TRN, somatosensory cortex, and many of their interconnections ^195,196^.

Mice were deeply anesthetized with isofluorane and decapitated. The brains were removed while submerged in cold (4 °C) oxygenated (95% O2, 5% CO2) slicing solution containing (in mM): 3.0 KCl, 1.25 NaH2PO4, 10.0 MgSO4, 0.5 CaCl2, 26.0 NaHCO3, 10.0 glucose and 234.0 sucrose. Brains were then mounted, using a cyanoacrylate adhesive (Krazy Glue), onto the stage of a vibrating tissue slicer (Leica VT1200S) and cut into thalamocortical slices (300 μm thick, 35° tilt from coronal). Slices were incubated for ∼1min in the cold sucrose-based slicing solution, then transferred to a holding chamber for 20 min with warm (32°C) oxygenated artificial cerebrospinal fluid (ACSF) containing (in mM): 126.0 NaCl, 3.0 KCl, 1.25 NaH2PO4, 1.0 MgSO4, 1.2 CaCl2, 26.0 NaHCO3, and 10.0 glucose. Subsequently, slices were allowed to equilibrate in ACSF for 60 min at room temperature before imaging or recording.

### Live imaging

After equilibrating, slices were placed in a submersion recording chamber and bathed continually (2-3ml/min) with warm (32°C) oxygenated ACSF (above). Images of live sections (300 μm), centered over the somatosensory sector of the thalamus (–1.4mm from bregma), were acquired using a Nikon upright microscope with 4x objective and Photometrics Prime BSI sCMOS camera. Epifluorescent and transmitted light (bright-field) images were obtained to characterize the opsin-fluorescent protein expression in corticothalamic cells and their axonal projections, particularly to the TRN.

Before recording, slices were examined to ensure that cortical expression was restricted to the targeted cortical areas and layers of interest. Slices with the brightest and sharpest cortical projections to somatosensory TRN were generally selected for recording. In experiments comparing the S1 L6 drive of central and edge TRN cells (Figures 1-3, S2, S3), slices with the brightest projections to the *medial edge* of TRN were used. In experiments characterizing CT synaptic inputs to TRN cells from S1 (Figures 1-4, S2-3), the slices were severed between VPm and TRN with a scalpel following initial imaging, isolating effects of CT inputs to TRN and preventing possible confounding excitation from thalamocortical nuclei.

### Whole-cell recording

TRN cells were targeted for recording by anatomical positions and tdT expression in the PV-Flp x Ai65 or PV-Cre x Ai14 mice. Whole cell recordings were made using DIC-IR visualization through a 40x water-immersion objective and patch pipettes containing potassium based internal solution (130 mM K-gluconate, 4 mM KCl, 2 mM NaCl, 10 mM HEPES, 0.2 mM EGTA, 4 mM ATP-Mg, 0.3 mM GTP-Tris, and 14 mM phosphocreatine-K; pH 7.25, ∼290 mOsm). Electrophysiological data were acquired and digitized at 20 kHz using Molecular Devices Multiclamp 700B and a Cambridge Electronic Design (CED) Power Power1401-3A AD interface running Signal v7 software. Patch pipette capacitances were neutralized, and series resistances (typically 10–25 MOhm) were compensated online (100% for current-clamp, 60%– 80% for voltage-clamp). Signals were low pass-filtered at 10 kHz (current-clamp) or 3 kHz (voltage-clamp) prior to digitizing. All voltages were corrected for a 14mV liquid junction potential.

Pharmacological agents (Figures 3, S5) were applied through the ACSF bathing solution, and effects were measured at steady-state, at least 10 minutes from the start of perfusion. During experiments involving paired comparisons of central and edge TRN cells, the order of recording for each cell type was counterbalanced.

### Optical stimulation of corticothalamic pathways

CT axons expressing opsins (ChR2 or ChrimsonR) were optically stimulated using either white (Mightex LCS-5500-03-22), 455nm (Mightex BLS-LCS-0455-03-22), or 617nm (Mightex BLS-LCS-0617-03-22) wavelength light-emitting diodes (LEDs) controlled by a Mightex (BLS-1000-2) or Open Ephys Cyclops drivers. For experiments using both opsins, the 455nm LED was used to activate ChR2, while the 617nm LED was used to selectively activate ChrimsonR. For experiments using only ChR2, the white LED was used. Stimulus light was collimated and reflected through a 40x water immersion objective, resulting in a 400 um diameter light spot that was directed at opsin-expressing CT terminals by centering the light spot over the recorded cell. LED intensities ranged from 0.37 mW to 25.7 mW (usually 11.75 – 25.7). In experiments testing functional convergence of CT pathways using ChR2 and ChrimsonR (Figures 6 and S7-S8), both 455nm and 617nm LED stimuli were delivered at 15.54 mW using a multi-wavelength beam combiner (Mightex LCS-BC25-0515). Most light stimuli were delivered as 10 Hz trains of 1ms flashes, with 10 second intertrain intervals.

### Immunohistochemistry

Following live imaging and recording (above), the 300 um thick slices were fixed for 1-3 days in 4% paraformaldehyde (in phosphate buffer, 4 °C) then cut into 50 um thick sections on a vibratome. The 50 um sections then underwent fluorescent immunostaining using parvalbumin antibodies to identify TRN neurons, and GFP and RFP antibodies to enhance signals of fluorescent proteins expressed in the corticothalamic neuronal pathways.

In brief, sections were washed 2 times in 0.1 M phosphate buffer containing 0.15 M NaCl, pH 7.4 (PBS) (10 min per wash), pre-incubated for 2 h at room temperature with a blocking solution (10% normal goat or horse serum, 2% Triton X-100, 0.1% Tween 20 in 0.1 M PB), then incubated with primary antibodies in the same solution for 3-5 days at 4 °C. After the primary incubation, sections were washed 2 times in PBS (10 min per wash), pre-incubated for 2 h in the blocking solution, incubated with a secondary antibody solution for 1-2 days at 4 °C, then washed 5 times in PB (10 min per wash). Sections were mounted and coverslipped (CitiFlour CFM-3 mounting media Cat# 17979-20), then imaged at 4x-20x on an Echo Revolution fluorescent microscope.

Primary antibodies (diluted to 1:1000) were: Rabbit Anti-Parvalbumin (Swant/PV27a, Monoclonal), Mouse Anti-Parvalbumin (Swant/PV235, Monoclonal), Mouse Anti-Green Fluorescent Protein (Millipore/MAB3580, Monoclonal), Chicken Anti-Green Fluorescent Protein (MilliporeSigma/AB16901, Polyclonal), Rabbit Anti-Red Fluorescent Protein (Rockland/600-401-379, Polyclonal), Rabbit Anti-DsRed (Takara/632496, Polyclonal), Goat Anti-mCherry (Origene/AB0040-200, Polyclonal).

Secondary antibodies (typically diluted to 1:300) were: Alexa Fluor Plus 405 Goat Anti-Mouse (Invitrogen/A48255), Alexa Fluor Plus 405 Goat Anti-Rabbit (Invitrogen A48254), Alexa Fluor Plus 405 Donkey Anti-Rabbit (Invitrogen/A48258), Alexa Fluor 488 Goat Anti-Mouse (Invitrogen/A11001), Alexa Fluor Plus 488 Donkey Anti-Mouse (Invitrogen/A32766), Alexa Fluor 488 Goat Anti-Rabbit (Invitrogen/A11008), Alexa Fluor 488 Donkey Anti-Chicken (Invitrogen A78948), Alexa Fluor 568 Goat Anti Mouse (Invitrogen/A11004), Alexa Fluor 568 Goat Anti-Rabbit 568 (Invitrogen/A11011), Alexa Fluor Plus 555 Donkey Anti-Goat (Invitrogen/A32816).

### Data analysis

Analyses of electrophysiological data were performed using CED Signal 7, Molecular Devices Clampfit 11 and Microsoft Excel. Analyses of anatomical data were performed using ImageJ, Microsoft Excel, and GraphPad Prism7-10.

To plot anatomical profiles of the corticothalamic projections to TRN, fluorescent intensities of the projections were measured by drawing a horizontal ROI line from the medial to the lateral edge of TRN, through the center of the projection. The vertical ROI line thickness was 93 um (Figures 1, S1). Likewise, TRN cell positions were defined according to their medial-lateral locations in the nucleus. HO edge cells were located within 20% of the distance across the TRN from either the medial or lateral boundary, whereas primary central cells were located in the central 60% of the nucleus^60^. The TRN boundaries were determined by visualizing fluorescent TRN neurons in parvalbumin-tdTomato reporter mice (above), or by post-hoc IHC staining for parvalbumin, or by identifying the white-matter tracts outlining TRN that are visible in bright-field images.

Intrinsic bursting in the TRN cells (Figure 3B) was measured by first adjusting the steady-state potential to –74mV with intracellular current, then injecting a 1 sec pulse of negative current (–150 to –300 pA) to hyperpolarize the membrane potential to approximately –94 mV (mean = –93.8 ± 0.23 SEM mV). Release from such hyperpolarization often evokes high frequency bursts of action potentials, driven by activation of T-type calcium currents (Figure 3)^23,59,101–103,105,106^. We measured spike counts during these “offset bursts” as the number of spikes in the 150ms after termination of the negative current^60^.

Repetitive spiking properties evoked by positive intracellular current steps (1 sec durations) were measured from baseline potentials of –74 and –84 mV (corresponding to tonic and burst mode for central TRN cells; Figure S5)^60^. Spike frequency adaptation (Figure S5) was quantified by calculating an adaptation ratio, which was the frequency of the first 2 action potentials evoked by the current step divided by the frequency of the last 2 action potentials during the step. For each cell, an average adaptation ratio was computed across all sweeps in which the frequency of the last 2 action potentials ranged between 20 and 60 Hz.

Synaptic responses to optical stimulation of CT axons were measured from TRN neurons recorded in whole-cell current clamp and voltage clamp. In current clamp, spike counts were tallied for the 100 ms periods following the onsets of the optical stimuli. In voltage clamp, the area or amplitude of an evoked EPSC was measured relative to the baseline current averaged over the 2ms before the onset of each light pulse. The areas of the EPSCs were measured over the 99ms immediately following the offset of the light pulse. Peak EPSC amplitudes were measured from the first 10ms following the offset of the light pulse. Values are average responses to 5-20 stimuli (typically 15).

EPSCs evoked by L6 CT stimuli were larger for central than for edge TRN cells when equal optical stimulus intensities were applied (Figures 2, S2, S3). To test whether these stronger synaptic inputs were necessary for the stronger synaptically-evoked spiking in central cells, optical intensities were lowered for the central cells in each central/edge pair to determine the optical intensity at which the central EPSC size matched that evoked in the edge cell (the edge responses were evoked at fixed optical stimulus intensities for these experiments; 11.8 mW for Figure 2, 23.5 mW for Figure 3). CT-evoked spiking in the paired central and edge cells were then compared using optical intensities that evoked the matching EPSC sizes. Generally, no single LED intensity evoked central EPSCs that exactly matched those in the paired edge cell. Thus, in most cases two LED intensities were used for the central cells, one that evoked EPSCs slightly stronger than those of the edge cell, and the other slightly weaker. The corresponding central cell spike counts evoked by these two intensities were averaged together, weighted by their relative differences in EPSC sizes from those of the edge cell.

### Statistical Analysis

Statistical comparisons were performed using GraphPad Prism7-10. Statistical tests used are indicated in the main text and figure legends. In plots of group data, center values are means and error bars show SEM. Statistical significance was defined as P < 0.05, unless otherwise noted.

## Acknowledgments

We thank Barry Connors for his continuous support, generosity, and insight. This study could not have been done without him. We also thank Tina Voelcker, Shane Crandall, Omar Ahmed, Linda Wadiche, and Jacques Wadiche for their many contributions. The work was supported by NIH R01 NS100016 and UAB Impact funds.

## Author Contributions

C.D.P., R.I.M.-G., and S.J.C. designed the experiments. C.D.P., R.I.M.-G., H.L., L.F.D., W.O.G. and S.J.C. conducted the experiments. C.D.P., H.L., L.F.D., and S.J.C. analyzed the results. C.D.P. and S.J.C. wrote the paper.

## Declaration of interests

The authors declare no competing interests.

## Supplemental information

Figures S1–S8

## Supplemental Figures

**Figure S1.**
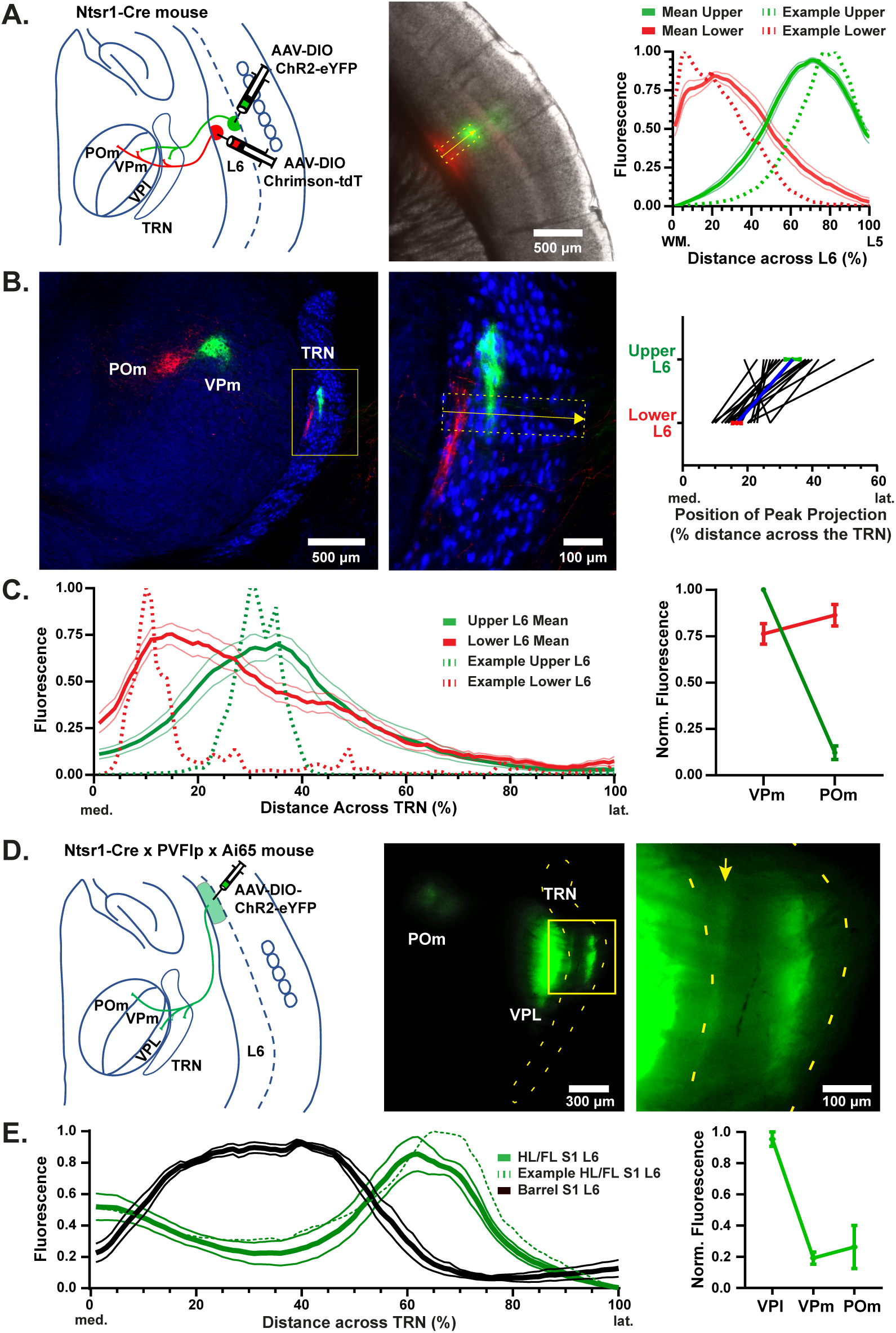
L6 CT inputs to separate zones of somatosensory TRN originate from cells in distinct regions and sublayers of S1. (A) **Left**, diagram of dual AAV injections in S1 L6, and corresponding CT projections to the thalamus. Small volumes (∼1/9 μl) of AAV2-DIO-ChR2-eYFP and AAV2-DIO-ChrimsonR-tdT were injected into upper or lower L6 of barrel cortex in Ntsr1-Cre mice. **Middle**, image of a live slice illustrating sublaminae-specific barrel cortical expression of ChR2-eYFP and ChrimsonR-tdT (gray brightfield image shows cortical laminae, including barrels in L4, superficial to the bright fluorescent signals). **Right**, fluorescence profiles illustrating patterns of cortical ChR2-eYFP and ChrimsonR-tdT expression for the example (dotted lines) and for a population of 18 mice with dual L6 injections (means ± SEMs, solid lines). Profiles for each mouse were obtained using ROIs lines drawn from the white matter to the L5/L6 boundary, intersecting both injection sites (like the yellow ROI drawn over the middle image; 162.5 μm wide ROI lines were used for cortical expression profiles). (B) **Left**, image of somatosensory TRN/thalamus from a 50 μm thick section (from the slice in panel A) following immunohistochemistry for YFP (green), tdT (red), and parvalbumin (blue) to identify locations of projections from upper and lower L6 relative to TRN/thalamic nuclei boundaries. **Middle**, magnified image of TRN showing CT fluorescent projections. ROI for measurement of fluorescent projection profile shown (yellow); ROI line thicknesses for TRN profiles were 93 um. **Right**, medial-lateral locations of the fluorescent profile peaks from upper and lower L6 CT projections to TRN. Lower L6 projections peaked more medially than upper L6 projections (16.7% vs 33.9%; P<0.0001, paired two-tailed *t*-test, n=18). (C) **Left**, fluorescence profiles illustrating patterns of L6 CT projections to TRN for the example from panel B (dotted lines) and for the population (means ± SEM, n = 18, solid lines). **Right**, relative fluorescence densities of CT projections to VPm and POm from upper (green) or lower (red) layer 6. Upper layer 6 always projected more strongly to VPm than POm (18/18 mice), while lower L6 only moderately favored POm over VPm (12/18 mice), consistent with known anatomy^10,80,83^. (D) **Left**, AAV-DIO-ChR2-eYFP was injected into L6 of the Hindlimb/Forelimb/Trunk regions of S1 in Ntsr1-Cre x PVFlp x Ai65 mice. **Middle**, image of a live slice centered over the thalamus illustrating eYFP-fluorescing L6 terminals in POm, VPl, and TRN. **Right**, magnified image showing dense terminal fluorescence in the lateral half of the central zone and a moderate band of fluorescence along the medial edge of TRN (arrow). TRN was outlined using tdT-expression in the PV-positive TRN cells (tdT expression was driven by Flp-mediated recombination in the PVFlp x Ai65 crosses). (E) **Left**, fluorescence profiles (mean ± SEM) comparing patterns of L6 CT projections to TRN from S1 Hindlimb/Forelimb/Trunk (solid green; n=7 mice) and S1 barrel cortex (black, n=9 mice). The dotted line is the profile from the example image in panel D. **Right**, relative fluorescent densities of L6 CT projections to the somatosensory TC nuclei from the non-trigeminal subfields of S1 (Hindlimb/Forelimb/Trunk). Projections favored VPl and avoided VPm, consistent with expectations^197^.

**Figure S2.**
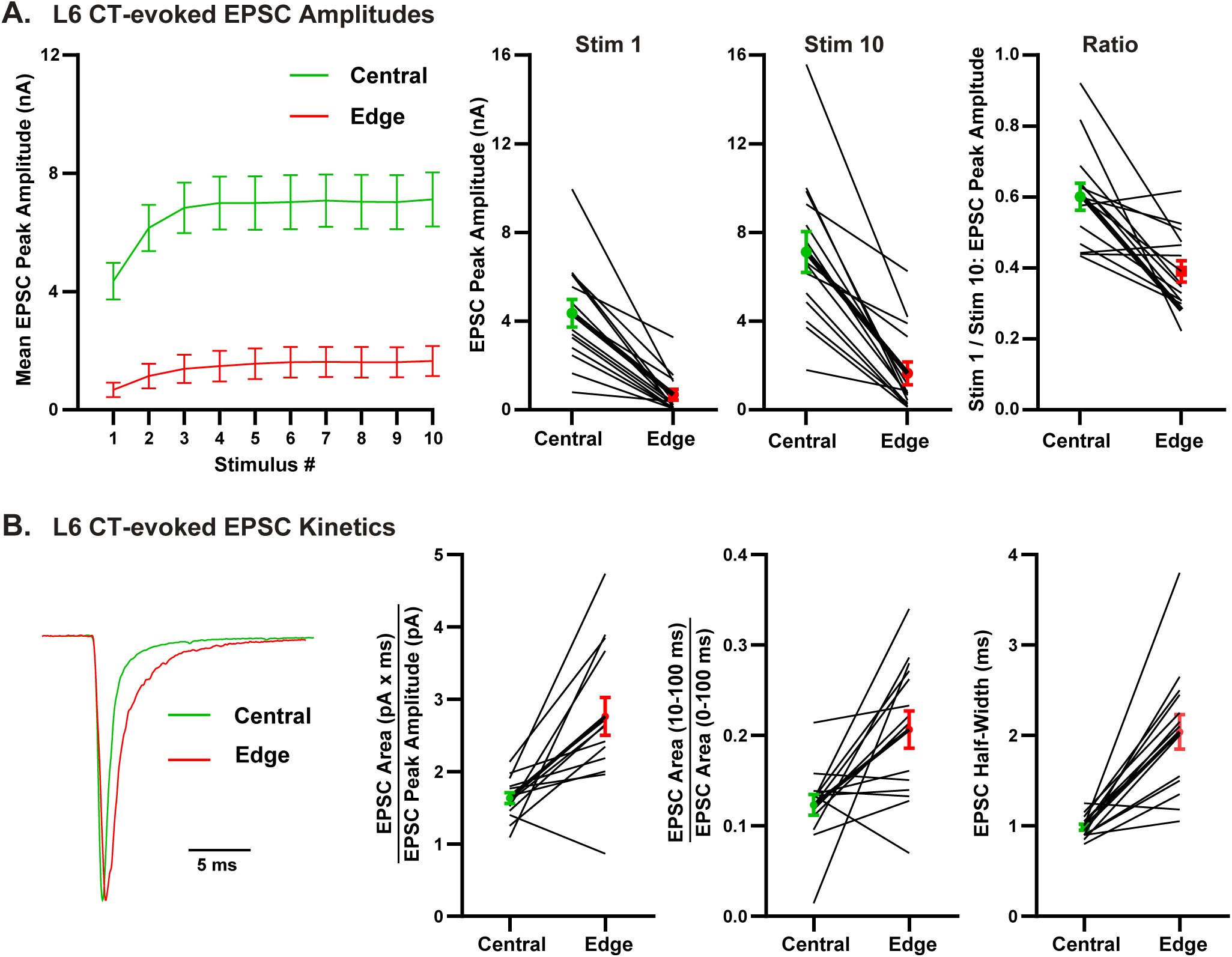
EPSCs evoked by L6 CT inputs in primary and HO TRN have distinct amplitudes and kinetics. (A) EPSC amplitudes and short-term plasticity. **Left**, mean peak EPSC amplitudes in central and edge TRN neurons evoked by 10 pulse optical train stimulation of S1 L6 CT axons (10 Hz trains, 1 ms pulses, 11.8 mW intensities; same n=14 central/edge cell pairs as Figure 2B-C). **Middle**, EPSC peaks were higher in central cells than edge cells for both the 1^st^ and 10^th^ stimuli of the trains (P<0.0002, paired two-tailed *t*-test, Bonferroni corrected for multiple comparisons; n=14 pairs from 12 mice). **Right**, the Stimulus 1/Stimulus 10 EPSC amplitude ratios were < 1.0 for all cells (and < 0.7 for 26/28 cells), consistent with robust short-term facilitation in all cell types. Facilitation was strongest in edge cells (P=0.001, paired two-tailed *t*-test). Thus, the characteristic magnitudes and short-term plasticity of L6 CT evoked responses in the distinct TRN subtypes were apparent whether those features were calculated from peak EPSC amplitudes (here) or from integrated charge (Figure 2). (B) EPSC kinetics. **Left**, EPSCs from a central and edge cell pair evoked by S1 L6 CT stimulation (10^th^ stimulus of the train from Figure 2B); the EPSCs were normalized to their peak amplitudes and aligned to response onsets. The edge cell EPSCs had longer durations. **Middle left**, EPSC Areas/EPSC Amplitudes were larger in edge cells (P=0.0008, paired two-tailed *t*-test). **Middle right**, the fractional areas in the late parts of the EPSCs were larger in edge cells (P=0.0067, paired two-tailed *t*-test). **Right**, EPSC widths were broader in edge cells (measured at half of the maximum amplitudes; P<0.0001, paired two-tailed *t*-test). All statistics in B: n=14 cells from n=12 mice. It remains to be determined if the cell type-specific differences in durations of these (optically-evoked) compound EPSCs result from longer duration unitary EPSCs or differences in synchrony of transmitter release among CT axons projecting to the two TRN cell subtypes.

**Figure S3.**
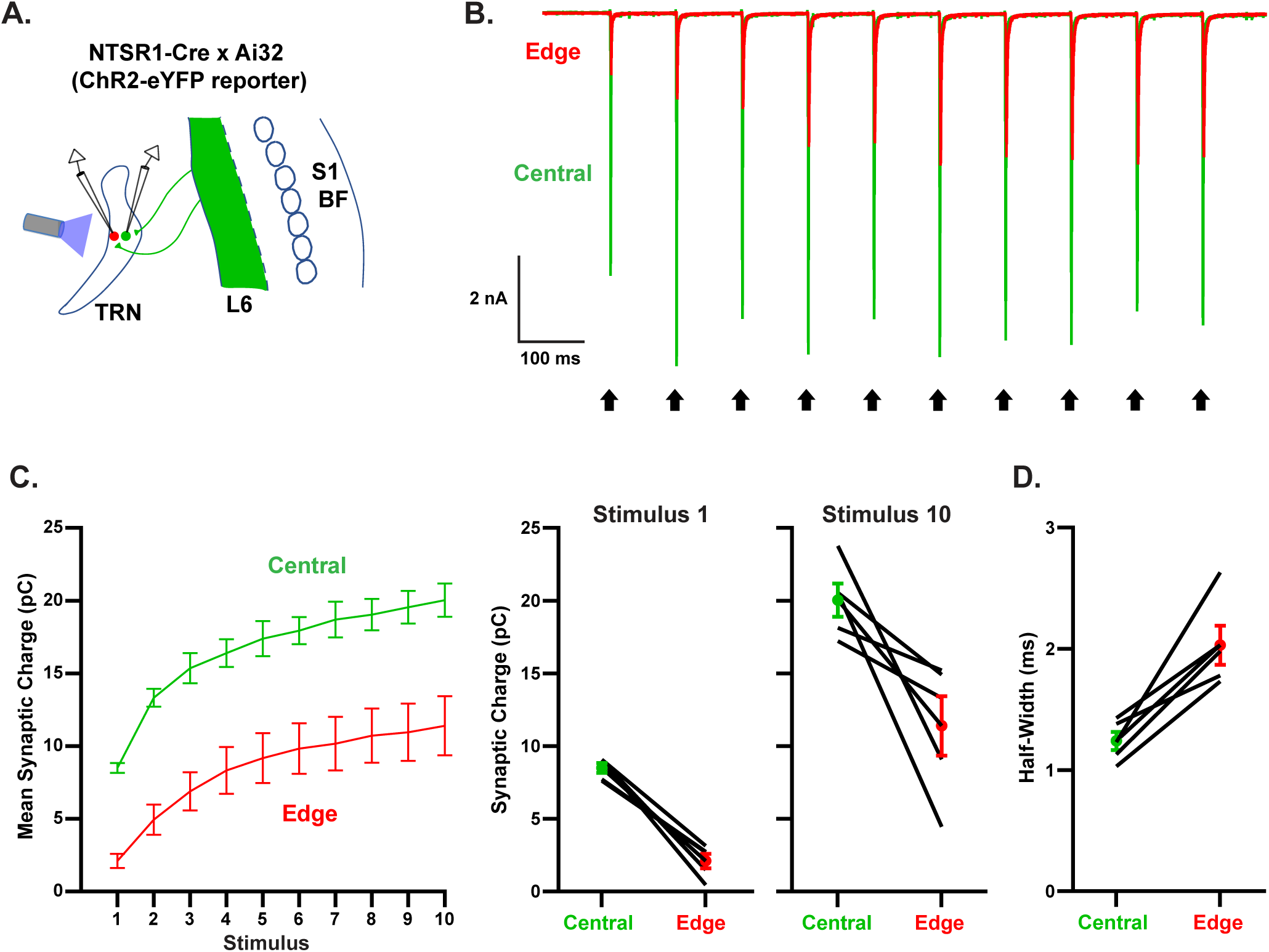
Properties of EPSCs evoked by L6 CT inputs in ChR2 reporter mice resemble those using viral ChR2 expression, confirming differences between primary and HO TRN. (A) Schematic of experimental setup. ChR2-eYFP was expressed in L6 CT neurons by crossing NTSR1-Cre mice with Cre-dependent ChR2-eYFP reporter mice (Ai32). Synaptic responses evoked by optical activation of the CT axons were compared in pairs of edge and central TRN cells (10 Hz pulse trains, 1 ms pulses, 23.5 mW). (B) Representative EPSCs from an edge and central cell pair. (C) **Left**, group plot of EPSC charge evoked by each stimulus in the train for the central and edge TRN neurons. EPSCs of both cell subtypes facilitated strongly, similar to responses evoked using viral ChR2 expression. Central cell EPSCs (green) were also stronger than those from edge cells (red), again similar to recordings obtained using viral methods. The integrated synaptic charges across stim 1-10 were (mean ± SEM, pC): Central cells = 166.2 ± 9.4, Edge cells = 84.54 ± 15.62 (P< 0.02, n = 5 cell pairs from 5 mice, paired two-tailed t-test). **Middle and right**, both the 1^st^ and 10^th^ stimuli of the trains evoked greater EPSCs in central cells than their paired edge cells (Stimulus 1, P*<*0.001; Stimulus 10, P<0.04, paired *t*-tests). (D) L6-evoked edge cell EPSCs had longer durations than those of central cells, again matching the pattern observed with viral expression of ChR2. The EPSCs half-widths were (mean ± SEM, ms): Central cells 1.24 ± 0.08, Edge cells: 2.03 ± 0.16 (P<0.01, n=5 pairs from 5 mice, paired *t*-test). EPSCs in the present figure were obtained from the same cells as those in Figure 3A-F and S5.

**Figure S4.**
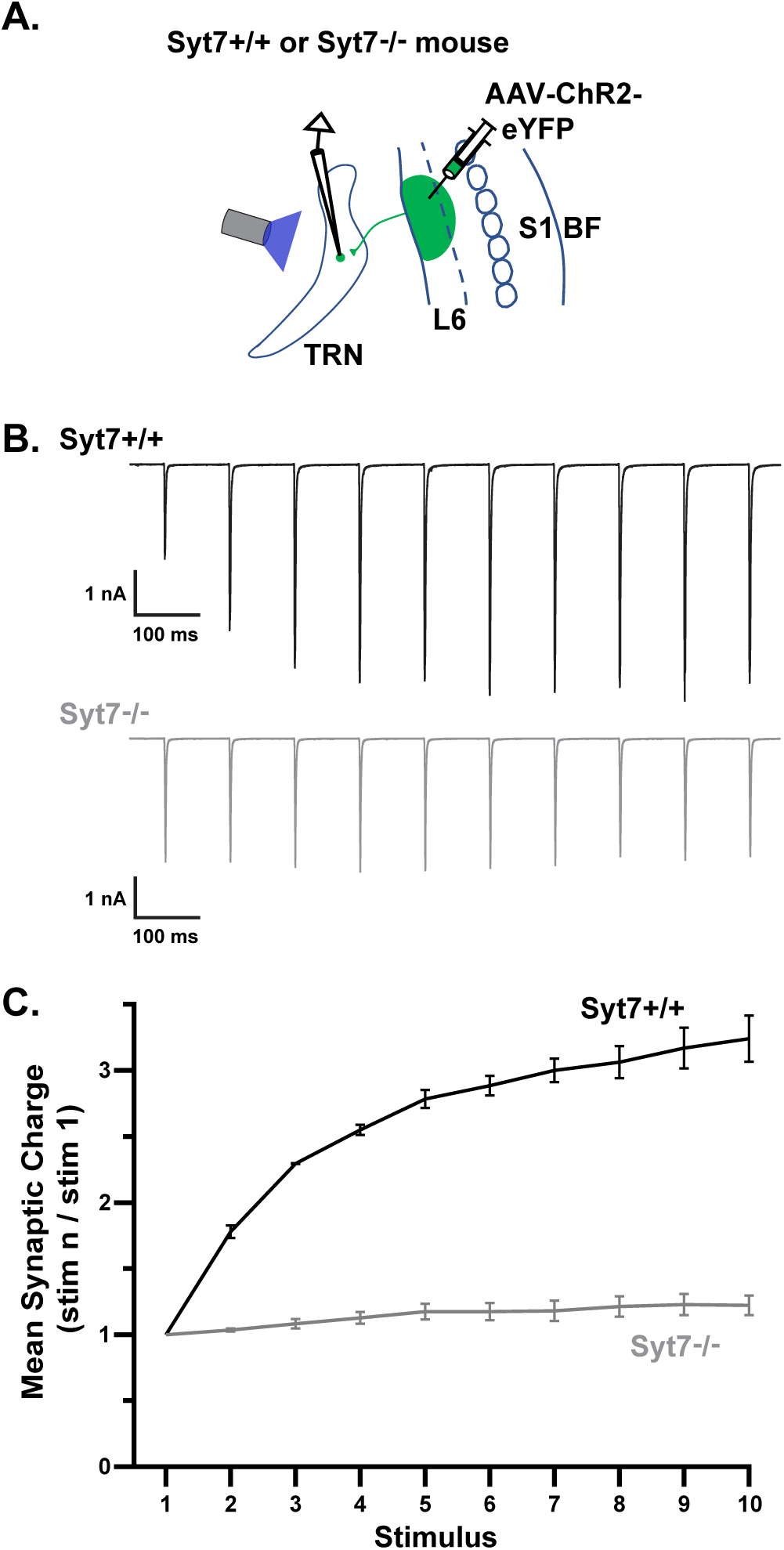
CT synaptic facilitation from S1 to TRN depends on Synaptotagmin 7. (A) Schematic of experiment. AAV-ChR2-EYFP was injected into infragranular S1 barrel cortex of Synaptotagmin 7 knockout mice or their wildtype littermates to induce ChR2-eYFP expression in CT cells. Subsequently, the EPSCs evoked by optical stimulation of CT axons were recorded in TRN and compared for the two genotypes. (B) Representative CT-evoked EPSCs of two central TRN cells, one from a wild-type mouse (top) and the other from a Syt7 KO (bottom). The responses from the Syt7 KO mouse did not facilitate. (C) Average (± SEM) short-term synaptic plasticity of CT-evoked EPSCs in TRN (n= 4 cells from 2 KO mice, n=3 cells from 2 WT mice; all primary central TRN). Short-term facilitation was nearly eliminated in Syt7 KO mice (Stim 10/1 EPSC charge ratios: WT = 3.24 ± 0.18, Syt7 KO = 1.22 ± 0.07, P= 0.0001, unpaired two-tailed *t*-test).

**Figure S5.**
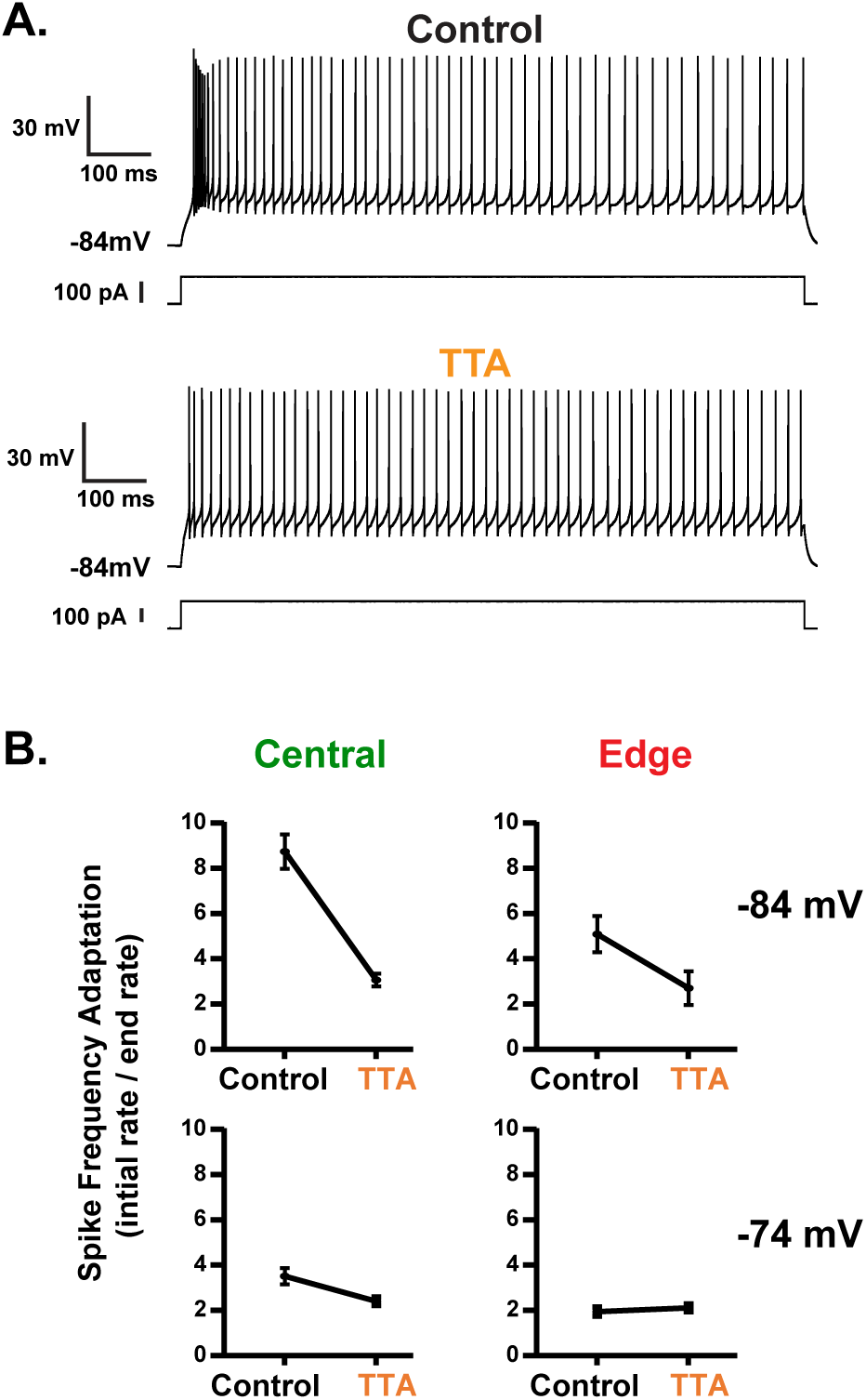
T-type calcium channels underlie differences in voltage-dependent spike frequency adaptation among TRN cell subtypes. (A) Example traces of central TRN cell spiking evoked by injecting a 125 pA positive current step before and during application of the selective T-type calcium channel blocker TTA-P2 (1 uM; –84 mV steady-state Vm). In control conditions, the depolarizing current evoked an initial high frequency burst, followed by a progressive reduction in spike rate (i.e., spike frequency adaptation). During TTA-P2, the initial burst was virtually eliminated, so the spike frequency adaptation was far less extreme. (B) The effect of steady state voltage and TTA-P2 on spike frequency adaptation. **Top left**, at –84 mV steady-state Vm, central cells in control ACSF responded as in the top trace of panel A, with an initial high frequency burst, followed by drastic reduction in frequency over the course of the current pulse. This resulted in large spike frequency adaptation ratios (= initial rate/end rate). Application of TTA-P2 blocked the initial bursting, leading to steadier overall rates and therefore lower spike frequency adaptation ratios (Adaptation ratios: Control = 8.74 ± 0.76; TTA-P2 = 3.06 ± 0.29). **Bottom left**, at –74 mV steady-state Vm, the central cells no longer burst strongly even in control ACSF (consistent with voltage-dependent inactivation of T-type calcium channels). Thus, there was only modest spike frequency adaptation. At this depolarized steady-state Vm, pharmacological blockade of T-type calcium channels had only a small effect (Adaptation ratios: Control = 3.51 ± 0.36, TTA-P2 = 2.40 ± 0.18). **Top right**, at –84 mV steady-state Vm, edge cell spike rates in control ACSF adapted, but not as strongly as rates of central cells, presumably due to less prominent T-type calcium currents. Consistent with this, TTA-P2 had weaker effects on adaptation in edge cells than on central cells (Adaptation ratios: Control = 5.09 ± 0.81, TTA-P2 = 2.70 ± 0.75). **Bottom right**, depolarizing the steady state Vm to –74 mV led to very weak spike frequency adaptation in the edge cells, and application of TTA-P2 had no clear effect (Adaptation ratios: Control = 1.95 ± 0.20, TTA-P2 = 2.11 ± 0.17). The recordings in B were made from the same 5 cell pairs (from 5 mice) as those in Figure 3A-F, with the exception of the upper left panel (Central, –84 mV), which included 4 cells recorded in the TTA-P2 condition.

**Figure S6.**
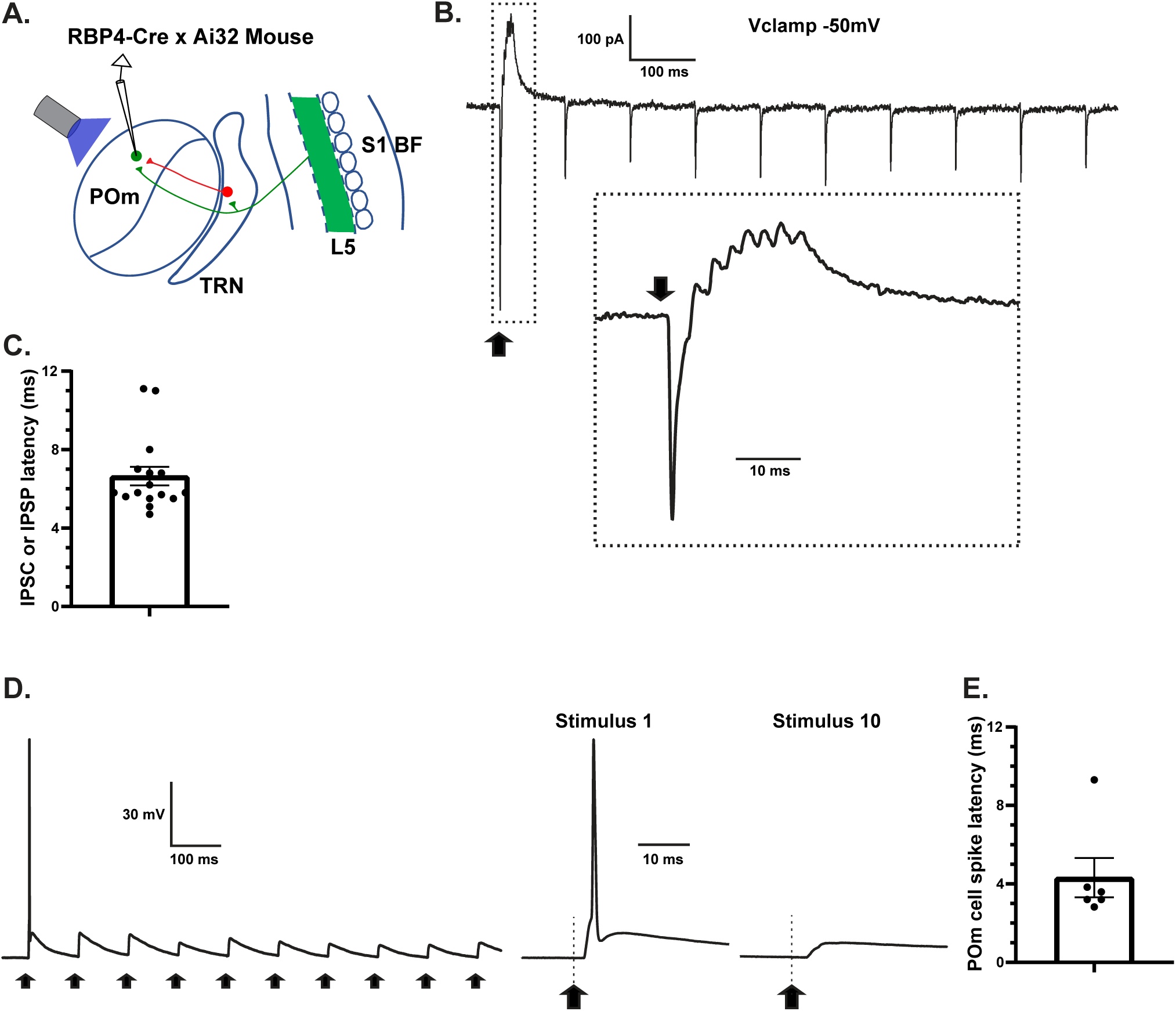
Activation of L5 CT projections evokes short-latency inhibition in POm, consistent with a disynaptic feedforward mechanism mediated through the TRN. (A) Schematic of experimental setup. ChR2-eYFP was expressed in L5 neurons by crossing RBP4-Cre mice with ChR2-eYFP reporter mice (Ai32). Synaptic responses evoked by optical train stimulation of L5 CT axons were recorded in POm thalamic neurons. (B) Example postsynaptic currents recorded from a POm cell clamped at –50 mV, a voltage between the GABA-A and glutamatergic reversal potentials (∼ –91 mV and +3 mV, respectively)^89^. **Inset**, compound PSC to the first stimulus in the train shown at a magnified time base. The response included an initial inward excitatory current, followed by a short latency outward inhibitory current. The arrow indicates the onset of the 1ms LED pulse. Both the inward and outward currents depressed strongly during the trains. The temporal dynamics of the responses were consistent with monosynaptic CT EPSCs, and disynaptic IPSCs. (C) Distributions of IPSC latencies recorded in POm cells in response to L5 CT stimulation (n=13 cells from 10 RBP4-Cre x Ai32 mice, and n=3 cells from 3 RBP4-Cre mice with AAV-DIO-ChR2). The majority of IPSC latencies (9/16) were < 6 ms. (D) **Left**, example trace from one of the POm cells that fired action potentials in response to L5 CT stimulation (6/16 POm cells had suprathreshold responses). **Right**, responses to the 1^st^ and 10^th^ stimuli in the train at a magnified time base. (E) Action potential latencies of POm cells in response to L5 CT stimulation (n=6). Based on the modest probability for spiking in POm (6/16 cells) and the latencies of POm spikes when they did occur, it is unlikely that the < 6 ms latency IPSCs observed in A-C were the result of a trisynaptic mechanism (L5 CT → POm → TRN → POm); the IPSCs are more readily explained by a disynaptic mechanism (L5 CT → TRN → POm).

**Figure S7.**
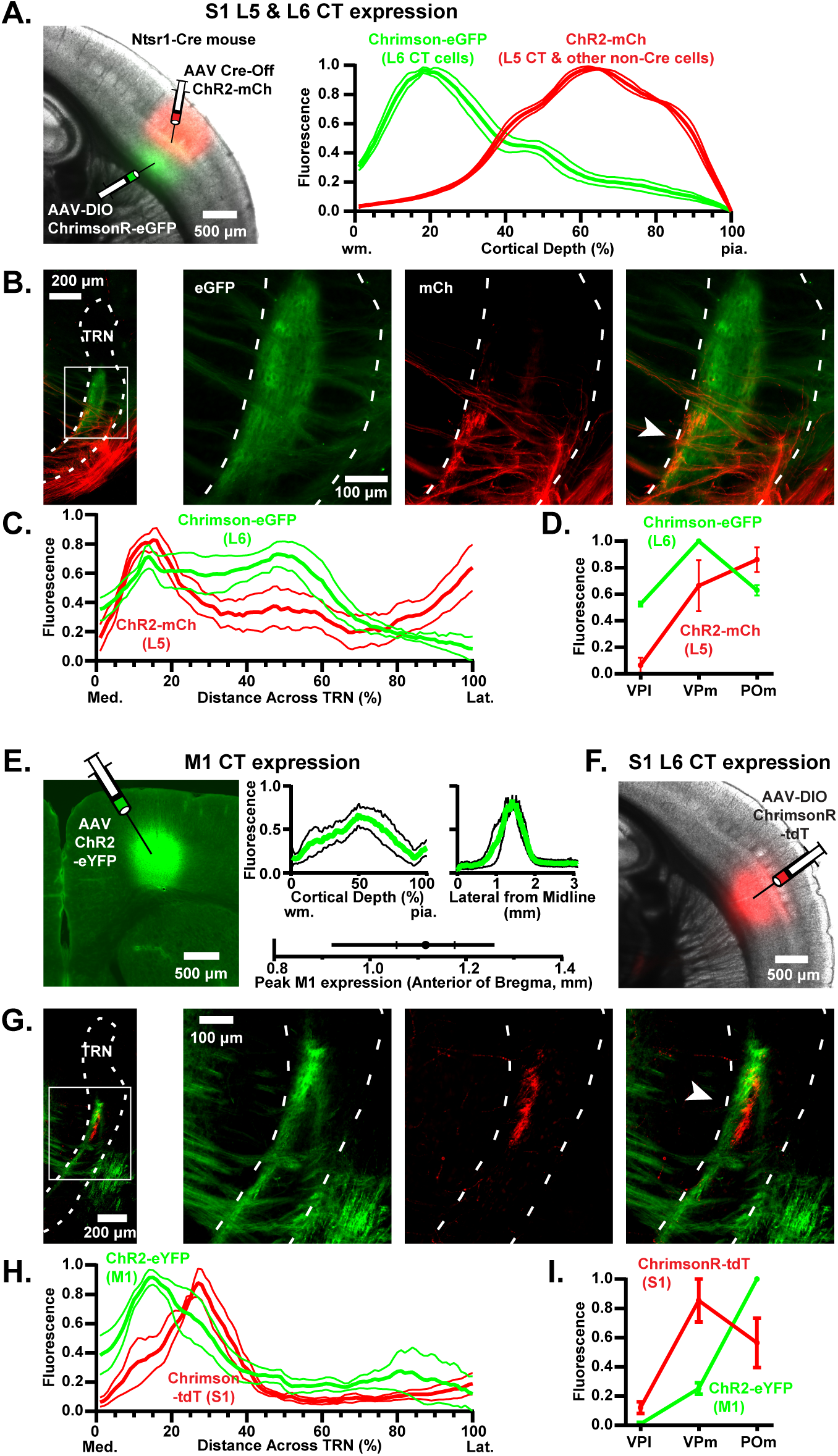
Projections from multiple CT systems converge on HO somatosensory TRN. (A-D) **Convergent input to TRN from L5 and L6 of S1.** (A) **Left**, image of a live slice centered on S1 barrel cortex of an Ntsr1-Cre mouse following infragranular injections of two AAV constructs. The first was a standard AAV-DIO construct, driving expression of ChrimsonR-eGFP in Cre-expressing L6 CT cells (green). The second was an AAV Cre-Off construct, driving ChR2-mCh expression in all neurons except those with Cre-recombinase (red)^137^; ChR2-mCh was expressed in L5 CT cells but excluded from L6 CT cells because the latter were Cre-positive. **Right**, mean fluorescence profiles (± SEMs; n=5 mice) showing ChrimsonR-eGFP expression in L6 and ChR2-mCh in L5 and above. (B) **Left**, example illustrating CT projections to the TRN from L6 (green) and L5 (red) following dual barrel cortical injections as in panel A. **Right**, magnified images of the framed region from the far left panel, with the L6/eGFP and L5/mCh signals shown separately and as an overlay to illustrate the region of overlapping terminals (arrow). (C) Group profiles of L6 (green) and L5 (red) S1 CT projections across the TRN (mean ± SEM; n=6 mice). Projections were roughly consistent with those from the single pathway experiments using Cre-on approaches in NTSR-Cre (Figs. 1, S1) and RBP4-Cre (Fig. 4) mice. Thus, the region of strongest L5/L6 terminal overlap in TRN was the HO medial edge zone. (D) Mean normalized fluorescent densities (± SEMs; n=6 mice) of L5 and L6 CT projections to the somatosensory thalamocortical nuclei. L6 CT cells projected most strongly to VPm and L5 most strongly to POm, again consistent with the single pathway experiments (Figs. 1, S1, 4)^66,68,89^. (E-I) **Convergent input to TRN from M1 and S1.** AAVs were injected into infragranular layers of motor and somatosensory cortices to drive expression of ChR2-eYFP in M1 CT cells (green) and ChrimsonR-tdT in S1 CT cells (red), then projections to thalamus/TRN were examined. The AAV constructs used in M1 were recombinase-independent for all M1/S1 convergence mice (AAV2-ChR2-EYFP; 4 mice), as were those in S1 for 2 of these mice (AAV2-ChrimsonR-tdT). For the other 2 mice, the AAVs in S1 were Cre-dependent (AAV2-DIO-ChrimsonR-tdT, NTSR1-Cre genotype), specifically targeting L6 CT cells. (E) **Left**, coronal slice illustrating expression of ChR2-eYFP (green) in infragranular vibrissa motor cortex. **Middle**, group plots showing locations (means ± SEMs, n=4 mice) of motor cortical ChR2-eYFP expression as a function of cortical depth, medial-lateral position, and anterior-posterior position (hash marks in anterior-posterior plot = SEM, line length = range). (F) Image of a live slice illustrating expression of ChrimsonR-tdT in L6 of S1 barrel cortex. (G) **Left**, example image illustrating CT projections to the TRN from M1 (green) and S1 (red). **Right**, enlarged images of the framed portion of the image from the left panel, with the M1/eYFP and S1/tdT signals shown separately and as an overlay to illustrate region of adjacent terminals (arrow). The example projection from M1 is the same as in Figure 5. (H) Group profiles of M1 (green) and S1 (red) CT projections across the TRN (mean ± SEM; n=4). The region of greatest M1/S1 terminal overlap is located around the border dividing the central and medial edge zones, approximately 20% of the distance from the medial to the lateral TRN boundaries. (I) Mean normalized fluorescent densities (± SEMs; n=4 mice) of M1 and S1 CT projections to the somatosensory thalamocortical nuclei. M1 projected most strongly to POm and S1 (barrel field) most strongly to VPm.

**Figure S8.**
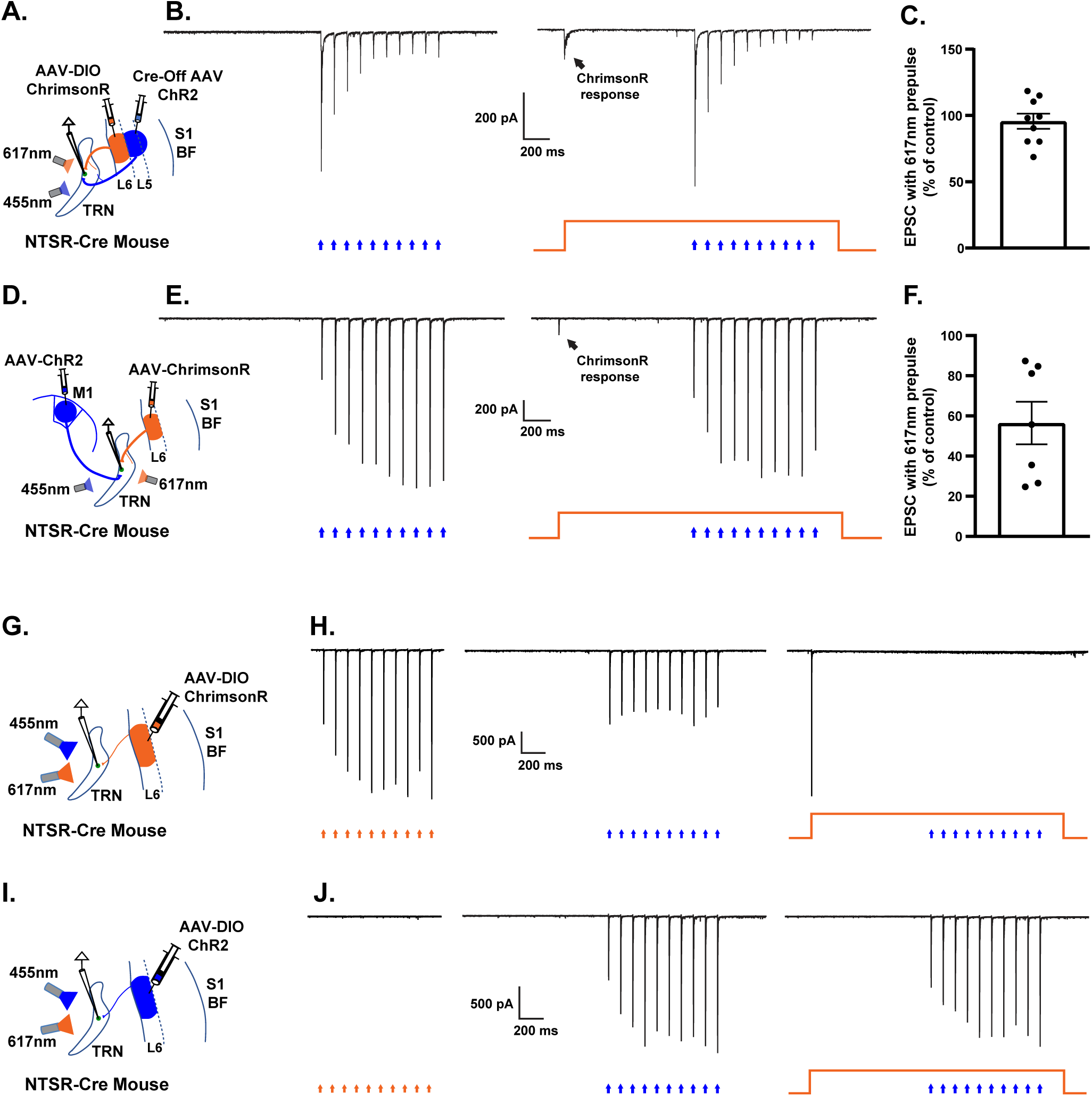
Separate groups of CT axonal projections that express ChR2 or ChrimsonR can be activated independently using 455 nm and 617 nm optical stimuli. (A) Experimental setup testing specificity of optical activation of ChR2-expressing L5 CT synaptic input. Using Ntsr1-Cre mice, ChrimsonR was expressed in Cre-positive L6 CT cells (red), and ChR2 in Cre-negative L5 CT cells (blue). The viral strategies are described in the Methods and are identical to those in Figures 6A-C & S7A-C. CT axons were stimulated with 455 nm and 617 nm LEDs at matching intensities, and EPSCs were recorded in medial TRN cells (where the L5 and L6 CT projections converged). (B) **Left**, EPSCs from a TRN cell evoked by stimulating CT axons with a train of 455nm light. **Right**, EPSCs from the same cell evoked by the same 455 nm stimuli delivered in the middle of a 2.1-second duration constant pulse of 617 nm light (the 455 nm train began 1 s from onset of the 617 nm pulse). The 617 nm stimulus prevented possible confounding synaptic input from ChrimsonR-expressing terminals that could potentially be activated by the 455 nm light train (as demonstrated in panel H below)^138^, thus isolating the synaptic input from ChR2-expressing terminals (i.e., L5 CT terminals). (C) The EPSCs evoked by the 455 nm LED train were not significantly attenuated by the 617 nm pre-pulse (P= 0.47, two-tailed t-test, n=9 cells). Recordings are from the same cells as those in Figure 6A-C. (D) Experimental setup testing specificity of optical activation of ChR2-expressing M1 CT inputs. ChR2 was expressed in M1 CT cells (blue) and ChrimsonR in S1 CT cells (red). The viral strategies are identical to Figures 6D-F and & S7E-H. CT axons were stimulated with matching intensity 455 nm and 617 nm LEDs, and EPSCs were recorded in medial TRN cells (where the M1 and S1 projections converged). (E) Same optical stimulation protocol as **B**. A 455 nm train stimulus was delivered either alone (**left**), or during a long duration 617 nm pulse designed to eliminate synaptic input caused by activation of ChrimsonR expressing terminals (**right**). (F) On average, the responses evoked by 455 nm trains in the presence of the 617nm pre-pulse was ∼ 56% of control (n=7 cells), indicating that there was a significant input to the recorded TRN neurons from ChR2-expressing M1 CT projections. The cells in E-F are the same as those in Figure 6D-F. (G) Experimental setup testing optical specificity for activating ChrimsonR-expressing CT terminals. Ntsr1-Cre mice were injected in L6 of S1 with AAV-DIO-ChrimsonR, driving expression of ChrimsonR in Cre-expressing L6 CT cells (red). CT axons were stimulated with 455 nm and 617 nm LEDs, and EPSCs were recorded in TRN cells. (H) **Left**, large EPSCs evoked by 617 nm stimulation of the ChrimsonR-expressing CT axons. **Middle**, smaller EPSCs evoked by 455 nm stimulation, consistent with the spectral properties of ChrimsonR^198^ (the 455 and 617 nm stimuli had matching intensities). Thus, 455 nm stimulation could excite ChrimsonR-expressing CT axons, potentially confounding interpretations without the use of proper controls. **Right**, the synaptic responses evoked by 455nm stimulation of ChrimsonR-expressing axons were eliminated by the 617 nm pre-pulse. This ensures that any responses evoked by 455 nm stimuli during such 617 pre-pulses were in fact mediated by ChR2-expressing axons (panels A-F and Figure 6). (I) Setup testing optical specificity for activating ChR2-expressing CT terminals of L6 CT cells (blue). CT axons were stimulated with 455 nm and 617 nm LEDs, and EPSCs were recorded in TRN cells. (J) **Left**, stimulating the ChR2-expressing CT axons with a 617 nm LED train evoked no synaptic responses in TRN. **Middle**, stimulating the same axons with a 455 nm train of the same intensity evoked robust EPSCs (middle), consistent with the spectral properties of ChR2^198,199^. **Right**, a 617 nm pre-pulse had no significant effect on EPSCs evoked by 455 nm stimulation of ChR2-expressing CT axons.

## Notes

### Competing Interest Statement

The authors have declared no competing interest.

### Summary of Updates

Language in the abstract was modified for clarity.

